# Understanding heterogeneous responses to T cell engagers: Binding characteristics and dosing thresholds determine cytotoxic efficacy

**DOI:** 10.64898/2026.02.03.703297

**Authors:** Tyler Simmons, Abhishek Kulkarni, Debasmita Paul, Freddys Rodriguez, Declan Dahlberg, Lakmal Rozumalski, Kamran Kaveh, Carston Wagner, Kevin Leder, Jasmine Foo

**Affiliations:** Therapy Modeling and Design Center, University of Minnesota; Department of Medicinal Chemistry, University of Minnesota; Department of Chemistry, University of Minnesota; Department of Biochemistry and Molecular Biology, Tulane University; Industrial Systems Engineering, University of Minnesota; School of Mathematics, University of Minnesota

## Abstract

Therapeutic efficacy of multivalent T cell engagers varies widely across individuals, but the basis for this heterogeneity remains poorly understood. Here, we integrate *in vitro* experiments of antitumor immune responses with a mechanistic modeling framework to investigate sources of response variability across T cell donors and TE constructs, focusing on a novel multivalent bispecific T cell engager currently in development. We identify parameter regimes that accurately recapitulate dose-response behaviors across T cell donors and doses, and perform cross-validation studies that demonstrate the model’s predictive accuracy. We find that variability in therapy efficacy is governed by the relationship between binding affinity and dose. When dose exceeds the binding affinity, responses are relatively robust across donors; when dose is below the binding affinity, responses are more donor dependent. At smaller doses, the TE-specific shape characteristics of the tumor-binding dose response, its steepness in particular, is a key marker of therapy efficacy. More generally, our integrated modeling and experimental framework offers insights and tools that are applicable to other bispecific T-cell engagers, and provides a quantitative foundation for the systematic, *in silico* optimization of TE design and dosing strategies.

## Introduction

In recent years, interest in T cell engagers (TEs) has grown rapidly. TEs are a class of cancer irnrnunotherapies that enhance a patient’s endogenous T cells’ ability to target cancer cells. Many bispecific TE molecules are constructed with two single-chain variable fragments (scFvs) connected by a flexible peptide linker: one scFv binds a tumor-associated antigen on the surface of cancer cells, while the other targets a T cell-associated molecule such as CD3. By physically bridging T cells and tumor cells, TEs stimulate T cell activation and subsequent tumor cell killing [1]. In contrast to autologous cell therapies that require costly expansion of patient immune cells *ex vivo*, TEs are ‘off-the-shelf’ agents that can be manufactured efficiently, engineered to optimize therapeutic efficacy, and are accessible to a broad range of patients. The use of TEs in cancer treatment is gaining momentum, with over 250 clinical trials involving T cell engagers reported to date, and more than a dozen TE therapies receiving regulatory approval in the US and Europe in recent years [2, 3, 4, 5, 6, 7, 8, 9].

However, the majority of TE therapy regulatory approvals have occurred in the setting of hematologic malignancies. Advancing these therapies in solid tumors has remained a challenge, as response have been highly heterogeneous. To date, only a handful TEs have gained FDA approval in a solid tumor setting, e.g. tarlatamab, which was approved as a second-line therapy for small cell lung cancer patients [11] and tebentafusp, for advanced uveal melanoma [12]. Challenges to TE therapeutics for solid tumors can be partially attributed to tumor antigen heterogeneity or scarcity as well as barriers to tumor infiltration, underscoring the need to employ engineering approaches to enhance and optimize both tumor antigen specificity and T cell avidity. Another challenge to TE therapy is systemic cytokine-related toxicity. Although TE therapy is generally associated with a lower incidence of cytokine release syndrome (CRS) than other immunotherapies, CRS accounted for 85% of TE-related adverse events in a recent study [13]. These toxicities may be further exacerbated in the setting of solid malignancies, where limited penetration and uptake in the tumor environment result in more free TE remaining in circulation and potentially triggering systemic CRS.

TE efficacy is governed by many processes spanning multiple scales, from stochastic molecular binding and immune-tumor cell crosslinking to tumor-immune evolutionary dynamics [14, 15, 16]. Current approaches to TE design and clinical strategies rely heavily on empirical trial-and-error, an inefficient process that can lead to failure of otherwise promising candidates. These challenges are further exacerbated by next-generation TEs, which introduce multivalent and modular molecular architectures that can improve molecular binding and activation [17, 18, 19, 20]. Here, multivalency refers to the presence of multiple binding arms for one or both target receptors (e.g., for CD3 on a T cell and tumor-associated antigen on a tumor cell). This enables a single TE to bind several receptors simultaneously, thereby increasing the effective binding strength and promoting T cell activation and cytotoxic efficacy. Multivalency can lead to sustained tumor cell killing at lower doses and antigen densities, potentially broadening their therapeutic window. Such innovations offer design flexibility but also create a vast combinatorial design space, underscoring the need for quantitative modeling tools that can identify the key factors driving tumor response, reduce the design space, and accelerate optimization of next-generation TEs.

Here we focus on chemically self-assembled nano-rings (CSANs), which are a novel multivalent TE constructed in a ring structure [17, 21, 18] built from multiple interchangeable monomers comprised of dihydrofolate reductase (DHFR) scaffolds linked to targeting ligands such as scFvs, fibronectins, affibodies, VHH domains and peptides that recognize cell-surface targets, commonly the CD3 complex of T cells and tumor-associated antigens, such as CD20, CD133, B7H3 (CD276), epithelial cell adhesion molecule (EpCAM), or epidermal growth factor receptor (EGFR) [22, 17, 23, 18]. These subunits self-assemble, with the help of bifunctional methotrexate (Bis-MTX) linkers, into a nano-scale, multivalent ring whose valency can be tuned [24, 25]. This multivalency enables improved targeting and linking of tumor and T cells, as well as the ability to address heterogeneous antigen expression within a tumor. Furthermore, these nano-rings can be rapidly disassembled with the approved antibiotic trimethoprim, providing a critical safety off-switch in the event of toxicity events such as the development of CRS. For both blood cancers and solid tumors, CSANs have demonstrated a strong *in vitro* efficacy as cancer immunotherapeutics [22, 17, 21, 23, 18]. Preliminary research indicates that CSAN-mediated T cell engagement and activation lead to robust tumor clearance. There have also been *in vivo* studies involving murine xenografts that also confirm the antitumor effectiveness of CSAN therapy [17, 18, 21, 23]. However, many *in vivo* studies of CSANs remain at the preclinical stages, with a strong focus on pharmacokinetics, tumor microenvironment infiltration, and patient safety during therapeutic intervention.

With initial preclinical *in vivo* and *in vitro* studies demonstrating the promise of CSAN therapies, the next steps require the development of guiding principles to optimize the design of such molecules. Computational modeling provides a quantitative framework for understanding how TE design choices and dosing strategies influence tumor-immune interactions and efficacy, across varying biological contexts such as antigen expression heterogeneity and T cell variability. Such a framework can be integrated with experimental data to optimize TE design and administration strategies, tuned to patient-specfic characteristics. Experimentally validated computational modeling frameworks can enable prioritization of the most promising TE variants and reduce reliance on preclinical animal studies, ultimately accelerating translation to clinic.

Mathematical modeling has increasingly been used to understand efficacy and guide development of TE therapies. Several mechanistic studies focus on the processes of tumor-immune complex formation and its dependence on target expression, binding affinity, and drug concentration ([26, 27]). Some works, such as those by [28] and [29], incorporate empirical data and make dosing predictions or simulate cell killing, while others like [30], [31], and [32] incorporate tumor growth modeling and biomarker identification. In [33], the authors examine cytokine release dynamics following TE therapy using a logic-based model. Broad overviews or tutorials for mathematical modeling of TE therapies are detailed in [34] and [35]. Although there is growing interest and substantial progress in using mathematical modeling to understand and guide TE therapy development, important gaps remain in linking mechanistic models of complex formation to tumor-immune population dynamics, incorporating inter-individual variability, and examining differences across TE constructs.

Here we develop an integrated modeling and experimental investigation of next-generation TE therapy using multivalent CSAN constructs. We quantify CSAN efficacy across multiple T cell donors and doses, and use these data to guide the development and parameterization of a population-level model of CSAN-driven tumor-immune dynamics. Through model reduction, parameter inference, and sensitivity analysis, we identify drivers of heterogneity in therapy response. This framework provides a quantitative foundation for the systematic, *in silico* optimization of CSAN design and dosing strategies, and offers insights and tools that are applicable more generally to other bispecific T-cell engagers.

## Results

### Experimental studies demonstrate dose-dependent tumor response to CSAN therapy *in vitro*

We experimentally evaluated the antitumor activity of two bispecific CSAN constructs, El-DHFR^2^ / *α*TCR-DHFR^2^ and DHFR^2^-E1/*α*TCR-DHFR^2^. Both CSAN constructs target the T cell receptor (TCR) on T cells and the EGFR on tumor cells, and were assessed for antitumor activity against a human epidermoid carcinoma tumor cell line (A431R). The CSAN constructs differ in orientation of the tumor-targeting ligands. In particular, distinct CSANs consisted of monomers with either an El-DHFR2 protein or a DHFR^2^-El protein (N-terminus to C-terminus). Here, El is a human tenth type III fibronectin that binds EGFR [36]. The two conformational structures have distinct binding affinities for EGFR, yielding dissociation constants of 60 nM and 120 nM, respectively, herein referred to as CSAN60 [21] and CSAN_120_. In contrast, the *α*TCR ligand binds TCR with a dissociation constant of approximately 1000 nM. A schematic of both CSAN constructs is provided in Figure 1. Across experiments, T cells were obtained from multiple donors. CSAN_60_ studies used T cells from three donors (D32, D41 and D42), while CSAN_120_ studies used T cells from two donors (D31, D35). For more details, see Supplementary Methods.

**Figure 1:**
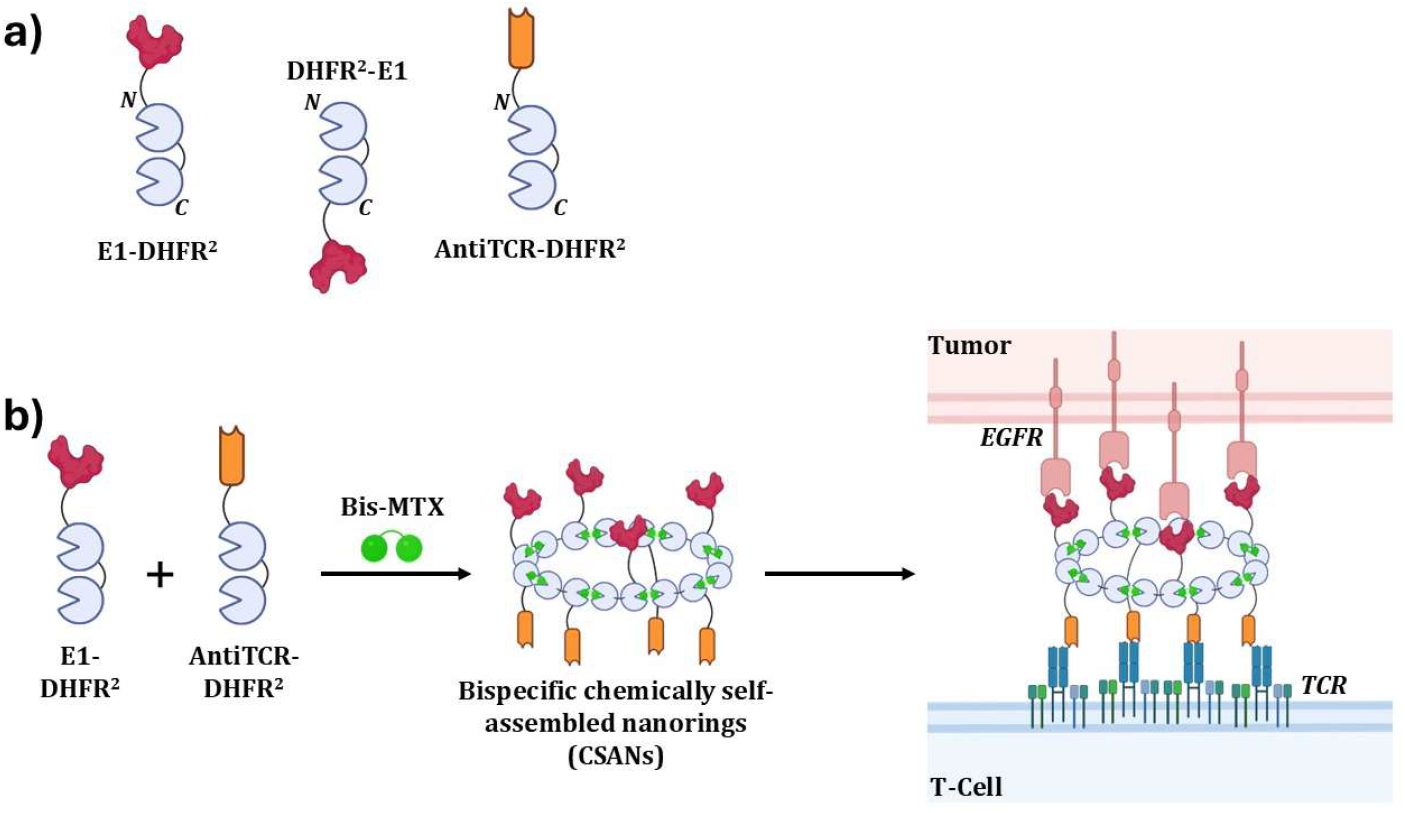
a) Representative diagram of the monospecific proteins used in each CSAN construct. b) An exemplary bispecific CSAN utilized in this work, consisting of El-DHFR^2^ monomer and antiTCR-DHFR^2^ monomer oligomerized via bis-methotrexate (Bis-MTX). This representative CSAN is an octomeric ring of equal valency for both targeting ligands, displayed in the trans-orientation, signifying distinct sides/direction for each ligand. The bispecific CSAN can induce a tumor cell: T-cell interaction leading to a T-cell-mediated tumor cell death.

#### Donor T cells alone provide negligible tumor killing of human epidermoid carcinoma in vitro

We first established the baseline growth kinetics of the A431R cancer cell line in the presence and absence of T cells from each of the five donors. For experiments with D32, D41, and D42, the initial T cell-to-tumor cell ratio (E: T) was 30,000:10,000 (3:1), while it was 50,000:10,000 (5:1) for D31 and D35. Counts of tumor cells only and tumor cells co-cultured with T cells were obtained using normalized fluorescence. Comparison of tumor growth dynamics in tumor-only cultures versus tumor /T cell co-cultures revealed low antitumor activity against A431R cells in four of the five T cell donor settings. In one donor T cell population (D41), tumor growth was statistically significantly reduced compared to the tumor-only control (see Supplementary Tumor Growth Statistics); however, the extent of tumor cell killing remained insufficient to reduce tumor burden.

#### Increased TE dose enhances efficacy against epidermoid carcinoma cells in vitro

To evaluate the concentration-dependent antitumor cell activity of CSANs in combination with donor-derived T cells, A431R cells were co-cultured with T cells from donors D32, D41, and D42 in the presence of CSAN_60_ at concentrations ranging from 400 nM to 0.39 nM (4-fold serial dilutions, performed in triplicate). Similarly, CSAN_120_ was tested with T cells from donors D31 and D35 across concentrations from 400 nM to 1.56 nM (2-fold serial dilutions, performed in duplicate). Normalized tumor fluorescence values for each replicate and condition are presented in Figure 3. These data demonstrate robust, dose-dependent antitumor efficacy of both CSANs, with potency varying across T-cell donors.

**Figure 2:**
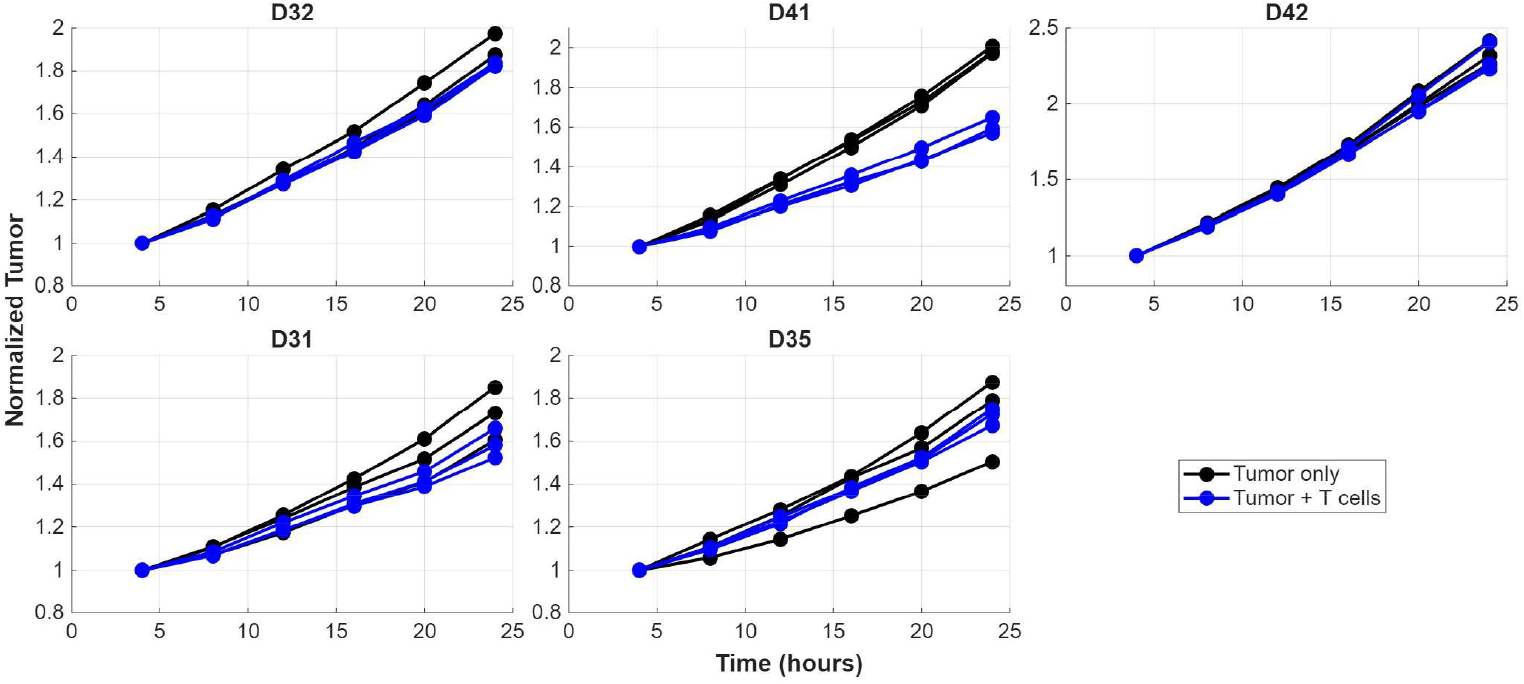
Baseline A431R human epidermoid carcinoma cell line growth (normalized fluorescence) in the presence (blue) and absence (black) of donor T cells. Three technical replicates are shown for each experiment.

**Figure 3:**
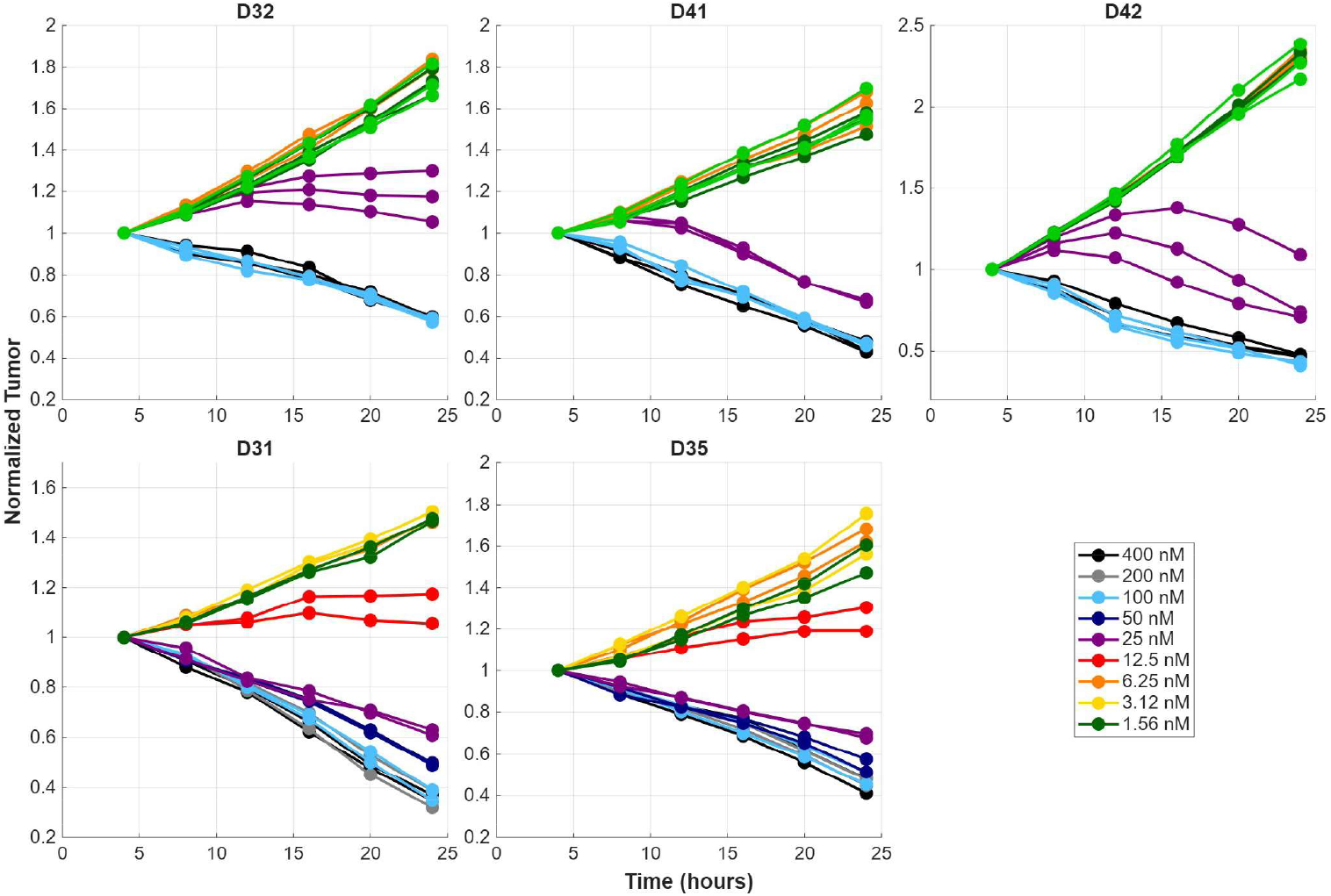
A431R human epidermoid carcinoma cell line growth (normalized fluorescence) in the presence of donor T cells and varying TE concentrations. (top row) Cancer cell population growth/shrinkage at varying CSAN_60_ concentrations with three donor T cell sets (D32, D41, and D42). Data has been capped at 24 hours for consistency across experiments; (bottom row) Cancer cell population growth/shrinkage at varying CSAN_120_ concentrations with two donor T cell sets (D31 and D35).

### Mathematical model of TE-mediated tumor-immune dynamics

To investigate the key mechanistic determinants of CSAN efficacy, we constructed a mathematical model of tumor-T cell dynamics in the presence of CSANs (Figure 4; eqs. (1)). The model describes the assembly of tumor-T cell complexes and tumor-CSAN-T cell trimers, followed by T cell-mediated tumor killing. Here, we use the trimer notation to indicate the tumor-immune complex formed through mediation of a CSAN, where synapse formation in this particular complex consists of three parts; tumor cell, T cell, and CSAN. This notation has been used previously in the literature for mathematical models of TEs [37, 26, 27]. All parameters of eqs. (1) are defined in Supplementary Table S1.

**Figure 4:**
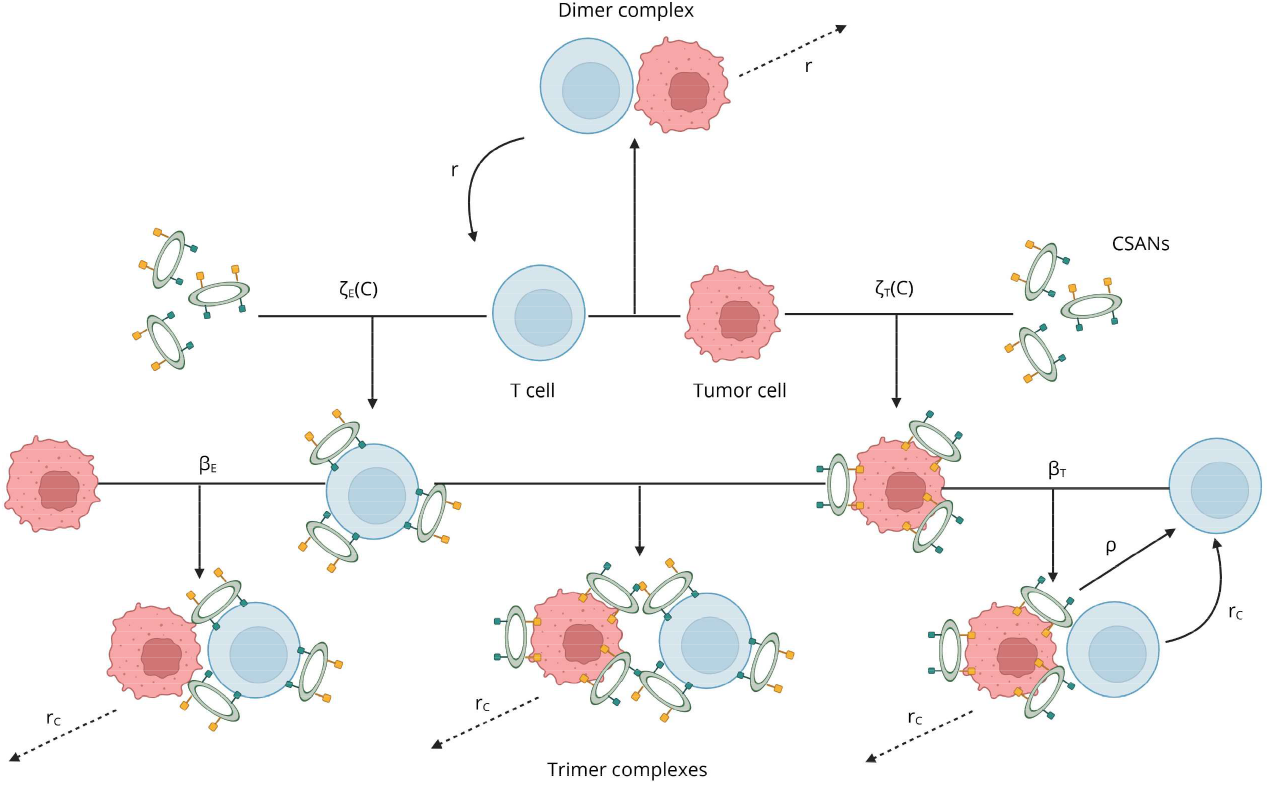
Schematic of the mathematical model of tumor-immune interactions in the presence of CSANs, see eqs. (1). Here, T cells may bind tumor cells, either independently or through CSAN mediation, forming complexes that may lead to T cell activation and tumor cell killing. Following target cell death (or disengagement), T cells are free to bind and form new complexes. Created in BioRender. Simmons, T. (2026) https://BioRender.com/ggm6ska

**Figure 5:**
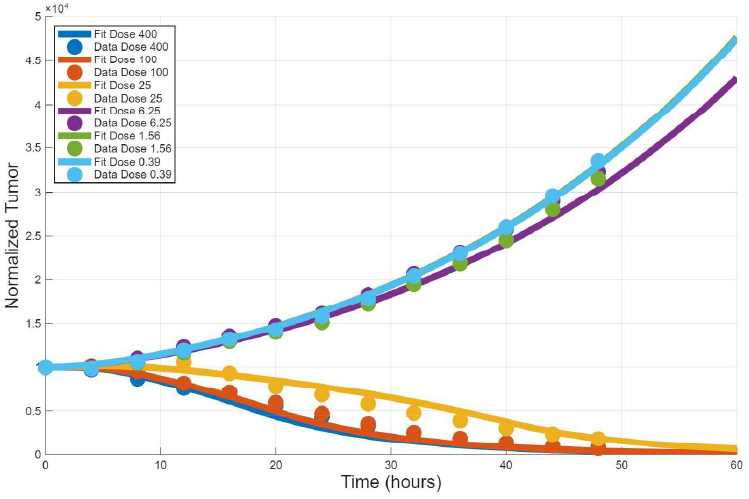
Calibration of model eqs. (1) to experimental CSAN response data. Tumor (A431R) and immune (D41) cells were mixed and treated with CSAN_60_. Fluorescence measurements of tumor cells were tracked over time. Data (markers) are the average normalized tumor fluorescence across dose-specific replicates at a given time. Solid lines depict the total tumor predicted by simulation of eqs. (1) using parameter estimates resulting from the fitting procedure to donor specific data. Model predictions are color-coded by simulated CSAN dose, corresponding to those used in experimentation.

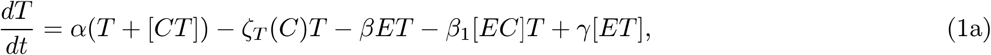

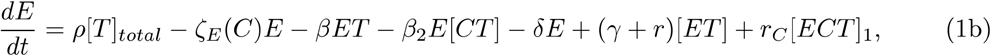

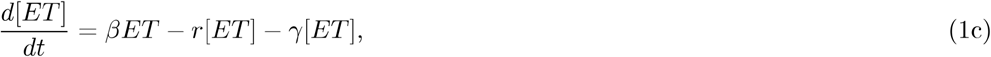

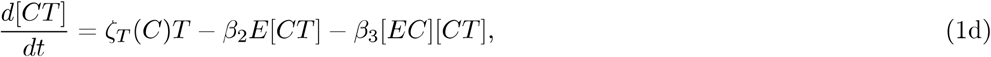

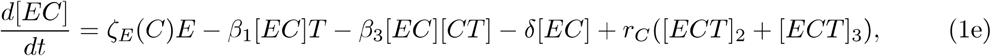

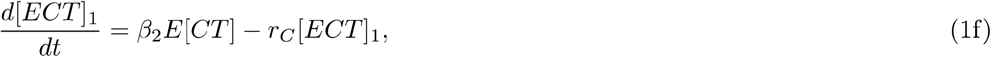

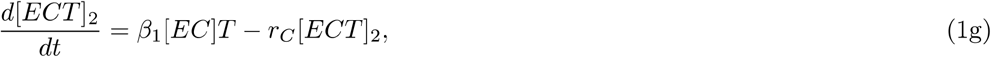

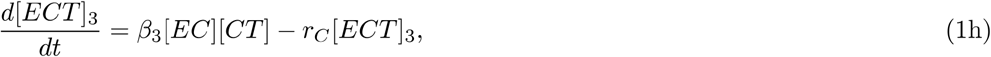

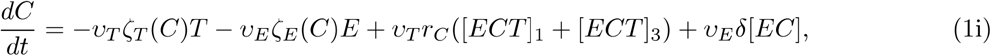

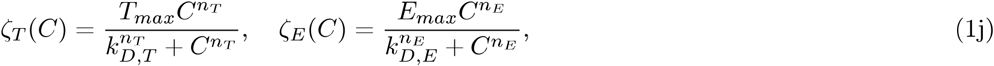

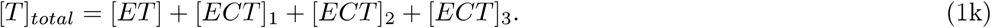

#### Tumor-T cell interactions in the absence of CSANs

Tumor cells, *T*, grow exponentially at rate *α* in the absence of T-cell-mediated killing (eq. (la)), as confirmed by tumor-only control curves in Figure 2. Even without CSANs, effector T cells, *E*, can recognize and bind to tumor cells, forming a tumor-T cell complex, at rate *β* (eq. (lb)). Bound tumor cells are killed with rate *r*, whereas the complex dissociates at rate *γ* (eq. (le)). The formation of tumor-T cell complexes may provide necessary signals to induce activation-derived proliferation, increasing the T cell population at a proportional rate given by the parameter *ρ*. For this stimulation, the tumor-immune complex is considered in [T]_*total*_, representing all tumor-T cell complexes, with and without CSANs. T cell exhaustion or death occurs at rate *α*.

#### CSAN binding and dissociation

As a simplifying assumption, we modeled each T cell and each tumor cell as existing in one of two discrete states: either CSAN-bound (e.g., [EC], [CT]) or CSAN-unbound *(E, T)*. In reality, the extent of CSAN engagement varies continuously, reflecting the high EGFR density on tumor cells and the TCR density on T cells. This continuum of receptor occupancy likely influences cellular behavior in a dose-dependent manner, resulting in a heterogeneous population of receptor-bound cells that interact to produce a complex landscape of antitumor activity. Resolution of the dynamics at this scale, as well as consideration of receptor-specific processes such as internalization, requires a stochastic, population dynamic model description, which is the subject of a future manuscript; here, we considered a simplified model to capture first-order dynamics as an initial step. In particular, we considered both tumor and effector cells to have two states: one in which binding to enough CSANs favors complex formation and immune cell killing, and one in which it does not.

The transition rates of tumor or effector cells into their respective CSAN-bound state takes the form of saturation-based dynamics given by Hill equations, denoted by ζ _*X*_(*C*), where the exponent nx dictates the steepness of this function for cell type *X* up to a maximum rate of *X*_*max*_ and where *k*_*D,X*_ represents the dissociation constant of CSANs targeting the same cell type (eqs. (ld), (le), (lj), (lj)). Following the binding of CSANs to either a tumor or T cell, the change in free CSAN concentration, measured in nM, is accounted for with the parameter *υ*_*X*_. Death or exhaustion of a CSAN-bounded T cell results in the now unbounded CSANs returning to the system (eq. (li)).

#### Trimer formation and tumor cell killing

Trimers are formed from interactions between T cells and tumor cells in which the T cell is CSAN-bound, the tumor cell is CSAN-bound, or both cells are CSAN-bound. To account for differences in free CSANs recycled back to the system upon death of a tumor cell from the trimer complex, we consider each of these processes separately in the model (eqs. (lf), (lg), (lh)). Binding of a tumor cell to a CSAN-bound T cell occurs at rate *β*_1_, binding of a CSAN-bound tumor cell to a T cell occurs at rate *β*_*2*_, and binding of a CSAN-bound tumor cell to a CSAN-bound T cell occurs at rate *β*_3_.

CSAN-mediated killing from a trimer complex occurs at rate *r*_*C*_. The recycling of CSANs back into the system following cell death depends on the underlying mechanism of cell death and the specific cell types involved. For example, effector T cell death only recycles CSANs back into the system if the T cell was part of the CSAN-bound compartment ([*EC*]). Likewise, tumor cell death recycles free CSANs only if the tumor cell in the trimer complex came from the CSAN-bound tumor compartment ([*CT*]). This principle holds for the recycling of cell types back into their appropriate compartments. To appropriately map the cell and CSAN recycling, the trimer complex must be considered as multiple compartments, *[ECT]*_1_, *[ECT]*_2_, and *[ECT]*_3_, where the first subset indicates the presence of CSANs on tumor cells only, the second subset for the presence of CSANs on effector T cells only, and the last subset for CSANs abundant on both cell types.

#### Additional model assumptions

We also assume that tumor cell proliferation occurs only in tumor cells not actively involved in a tumor-T cell or trimer complex. It is assumed that any daughter cell following division will not have enough CSANs bound to be considered in the CSAN-bound state, so new cells are considered unbound by CSANs. Additionally, we assume that due to CSAN mediation, trimer complexes are held sufficiently tight to enable appropriate signaling and cellular crosstalk between T cells and tumor cells to trigger tumor cell death prior to trimer dissociation.

##### Mathematical model captures dose-dependent tumor cell response to TE therapy

Tumor fluorescence measurements were fit to the mathematical model (eqs. (1)). Parameter fitting was done in a stepwise manner, where experimental data sets tracking tumor and immune dynamics in the absence of CSANs were used to parametrize donor-specific tumor growth rates, T cell exhaustion and death rates, tumor-T cell formation and dissociation rates, and the rates of T cell cytotoxic killing from the tumor-immune complex. These parameter estimates were fixed as constants within eqs. (1) for each T-cell donor. We calibrated the remaining parameters for each set of CSAN response experiments, identifying one parameter set to model data across all doses. An example model fit (D41 in the presence of CSAN_60_) is provided below, demonstrating that eqs. (1) recapitulated CSAN temporal dose-response dynamics well, across all doses. Details of the fitting procedure are provided in Supplementary Model Fitting.

#### Example model dynamics for donor D41

Figure 6 shows model predictions, using parameters calibrated to donor D41 in the presence of CSAN50, for unbounded tumor and effector T cells, the tumor-immune complex formed without CSANs, CSAN-bound tumor and effector T cells, all three trimer complexes, and the free CSAN concentration over time. Immediately, due to the large number of T cells present, tumor cells and T cells bound together to form the tumor-immune complex. With limited killing in this state, eventual complex dissociation resulted in the recycling of cells back to their individual populations. Large CSAN doses were then able to transition each cell type to their respective CSAN-bound state. These CSAN-bound cell types then started forming one of the three variants of trimer complexes. Comparing the magnitudes of each trimer complex, T cells bound to CSAN-bound tumor cells were the most frequent trimer formed. For larger CSAN doses, the total number of trimers formed, as well as the speed at which they were formed, were sufficient to promote tumor control. At lower doses, trimer formation was inadequate, resulting in tumor growth. Regardless of the CSAN dose, a limited change in the free CSAN concentration was observed.

**Figure 6:**
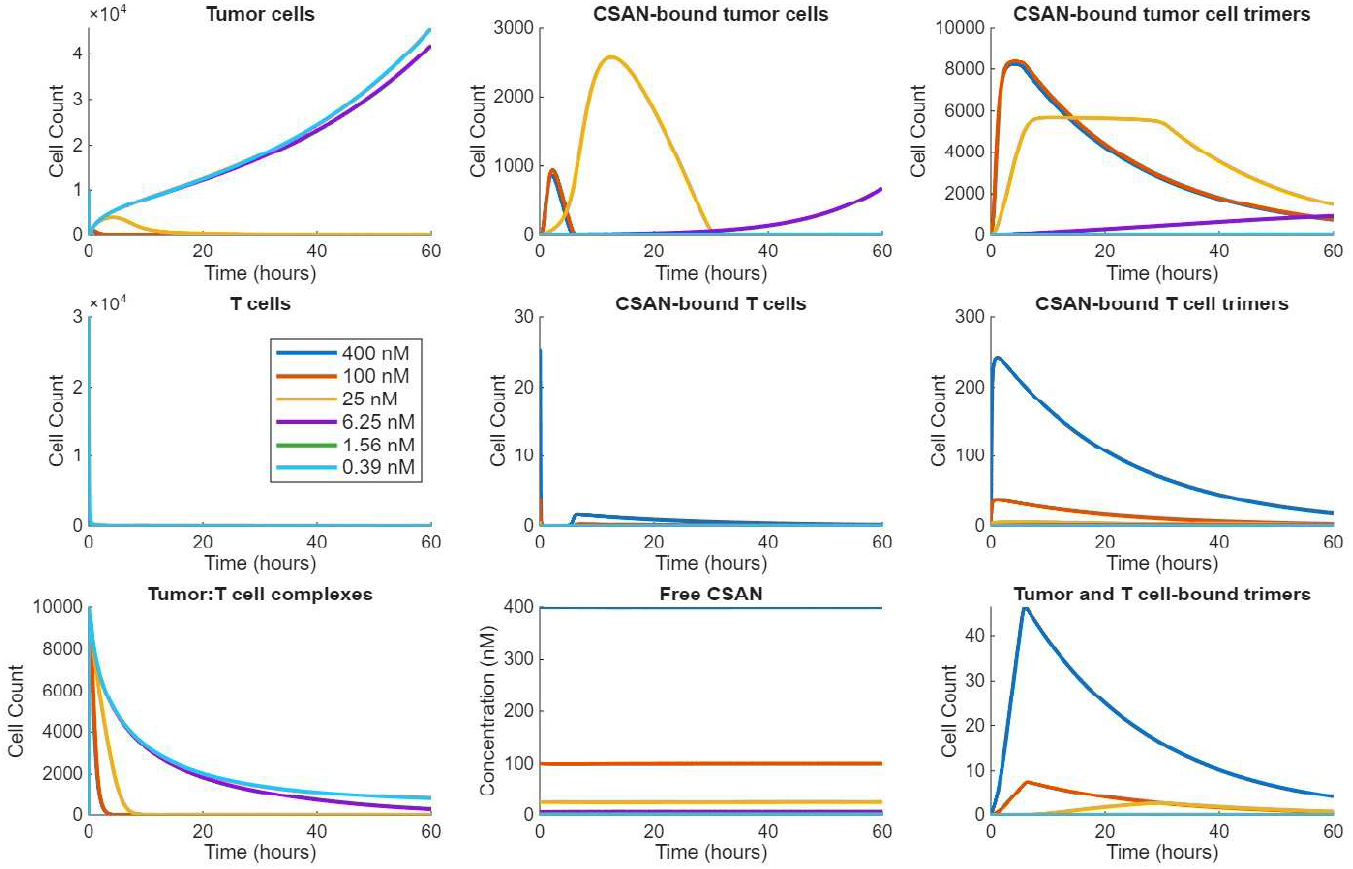
Simulated state variable trajectories of an example (D41) donor-specific parameterized model. State variables include: (top row) tumor cells, CSAN-bound tumor cells, and trimers consisting of CSAN-bound tumor cells with T cells, (middle row) T cells, CSAN-bound T cells, and trimers consisting of CSAN-bound T cells with tumor cells, (bottom row) tumor immune complexes without CSAN mediation, the free CSAN concentration (nM), and trimers consisting of CSAN-bound tumor cells and CSAN-bound T cells. Trajectory colors correspond to the same CSAN doses across all panels.

Our mathematical model, which describes TE binding, cellular interactions, and tumor-immune population dynamics, accurately captures the dose-dependent tumor response over time. It also provides deeper insight into the underlying dynamics of key species (e.g. T cells in bound and unbound states) driving the temporal evolution of tumor response. In the following sections, we will leverage this model to interpret CSAN therapy response using an integrated computational-experimental approach, incorporating key simplifications driven by experimental and modeling observations. More broadly, this approach is widely applicable and provides a general strategy for understanding and optimizing TE therapies.

### Data-guided model refinement

The full mechanistic model accurately recapitulated tumor CSAN response dynamics, capturing both temporal evolution and CSAN dose dependence. However, its structural complexity posed challenges to parameter estimation and identifiability, and resulting parameter fits exhibited limited consistency across donor-specific estimates. We then conducted sensitivity analyses and systematic model reduction to obtain a streamlined model representation that preserves accuracy while improving interpretability. For example, preliminary data (Figure 2) demonstrated negligible tumor killing by T cells without CSAN mediation and minimal T cell death/exhaustion during the time course; such interactions were removed in the reduced model.

#### Trimer formation is driven by CSAN-tumor binding interactions

Figure 6 demonstrates that CSAN-bound tumor cell abundance is consistently, significantly higher than CSAN-bound effector cell levels, indicating that trimers are primarily formed through the interaction of CSAN-bound tumor cells with a free effector cell, rather than the alternative pathway of a CSAN-bound effector cell with a free tumor cell. This is consistent with CSAN binding affinities towards T cells, which have a *k*_*D*_ of 1000 nM, much larger than towards the tumor cell. *Moreover, this behavior is consistent across all model parametrizations throughout our dataset, across donors and CSAN constructs*. This suggests a simplification of model eqs. (1) by removing CSAN-T cell binding events (eq. (lj)) and the CSAN-bound T cell compartment (eq. (Id)) as well as two of the trimer compartments with CSAN-bound T cells present (eqs. (lg) and (lh)).

Furthermore, as shown in Figure 6, the free CSAN concentration, governed by eq. (li), is relatively stable despite the binding of CSANs to cells and the recycling of CSANs after tumor cell destruction via T cell cytotoxicity. For a dosing range of 400 to 0.39 nM, given in a total volume of 200 *µL*, there are approximately 5 × 10 ^10^ - 5 × 10^13^ CSANs. There are roughly 6 × 10^6^ EGFR receptors per cell on A431R tumor cells and approximately 40 − 100 thousand TCR receptors on each T cell, resulting in approximately 6 × 10 ^10^ total binding sites within the experiment (when considering cell population counts). As the maximum number of receptors that could sequester CSANs is negligible relative to the majority of CSAN doses, we considered the CSAN concentration as a constant throughout the time course and removed eq. (Ii) and its parameters. We therefore fixed the CSAN concentration value to the initial dosage in eq. (lj), so that the transition from a tumor cell into the CSAN-bound state is dependent on the initial dose.

Following these simplifications, we introduced a reduced model that incorporates trimer formation via only the CSAN-bound tumor cell intermediary, shown in Figure 7 and in eqs. (2). Due to model simplifications, consisting of the elimination of various state variables and parameter values, parameter notation has been slightly altered. Notable differences include only one remaining tumor-immune binding rate, denoted *β*, and the CSAN-bound state transition rate of tumors, which is a function of initial CSAN dosage, *ζ* (*C*_0_). Additionally, the CSAN dissociation constant within *ζ* (*C*_0_) is simplified to *k*_*D*_ rather than *k*_*D,T*_. The parameters of the reduced model equations are described below in Table 1. To reduce the computational expense for parameter fitting, we nondimensionalized the model with respect to the tumor growth rate, *α*. This nondimensionalization is provided in detail in Supplementary Nondimensionalization.

**Table 1:** Parameter definitions with units, before nondimensionalization, for simplified model (eqs. (2)). Complexes consist of one tumor cell and one T cell. This one-to-one ratio eliminates the need for unitary conversion factors for the binding (complex association) and killing (complex dissociation) rates.

| Parameter (units) | Description |
| --- | --- |
| $\alpha$ ($hour^{-1}$) | Exponential tumor growth rate |
| $k_D$ ($nM$) | CSAN:tumor cell dissociation constant |
| $T_{max}$ ($hour^{-1}$) | Maximum transition rate of tumor cells to CSAN-bound state |
| $r_C$ ($cells \cdot (complex \cdot hour)^{-1}$) | CSAN-mediated tumor cell death rate |
| $n_T$ ($dimensionless$) | Hill coefficient of tumor cell transition to CSAN-bound state |
| $\beta$ ($(hour \cdot cell)^{-1}$) | CSAN-bounded tumor and T cell binding rate |
| $\rho$ ($cells \cdot (complex \cdot hour)^{-1}$) | Trimer induced stimulation and proliferation rate |

**Figure 7:**
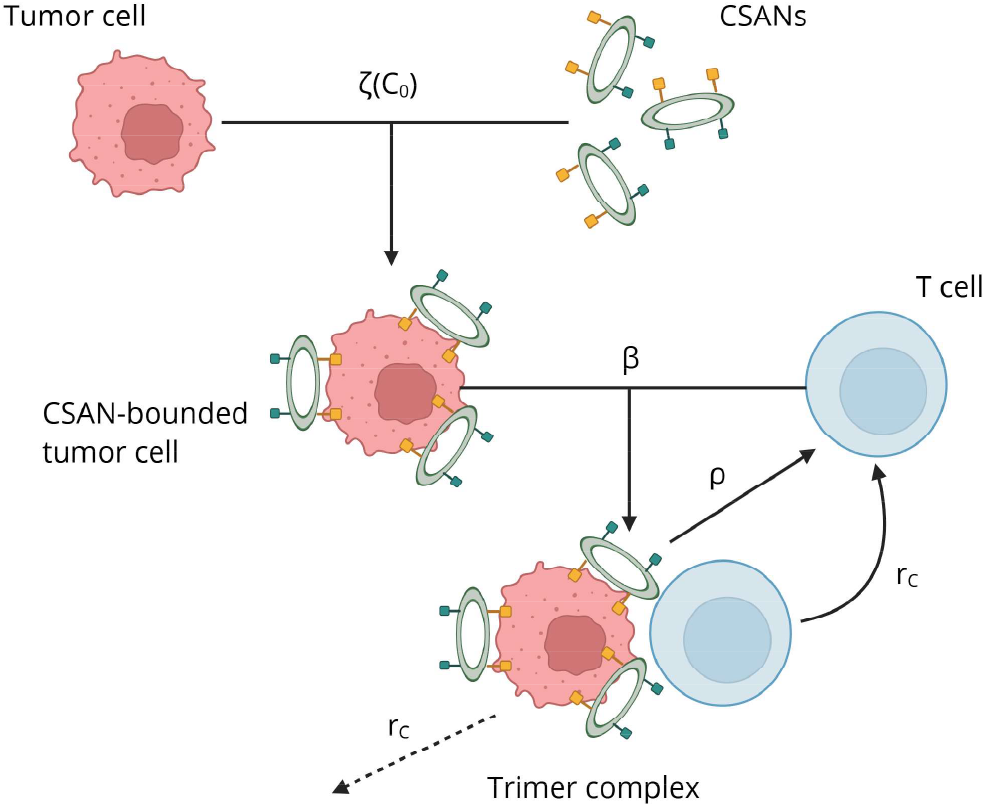
Schematic of simplified model equations, as described in Section Mathematical Model. CSAN-bound tumor cells are formed at rate *ζ* (*C*_0_), dependent upon the initial dose of CSAN, *C*_0_. CSAN-bound tumor cells my bind to effector T cells at rate *β* to form trimer complexes. In these complexes, T cell killing occurs at rate *r*_*C*_. T cells also experience population expansion proportional to trimer formation, denoted *ρ*. Created in BioRender. Simmons, T. (2026) https://BioRender.com/zfelyvt

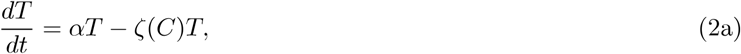

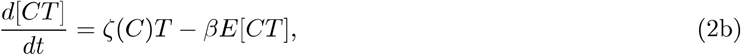

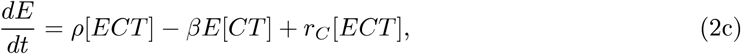

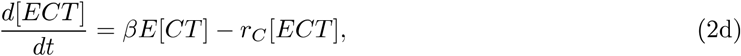

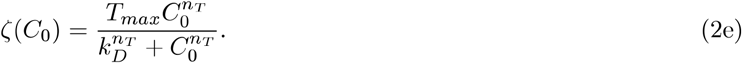

##### Reduced model accurately captures tumor response dynamics

The fitting procedure (see Supplementary Model Fitting) was performed on the nondimensionalized form of eqs. (2) (eqs.S10) for each dataset. The resulting simulation predictions and model fits for each donor are shown in Figure 8, demonstrating that the reduced model accurately recapitulates experimental dynamics.

**Figure 8:**
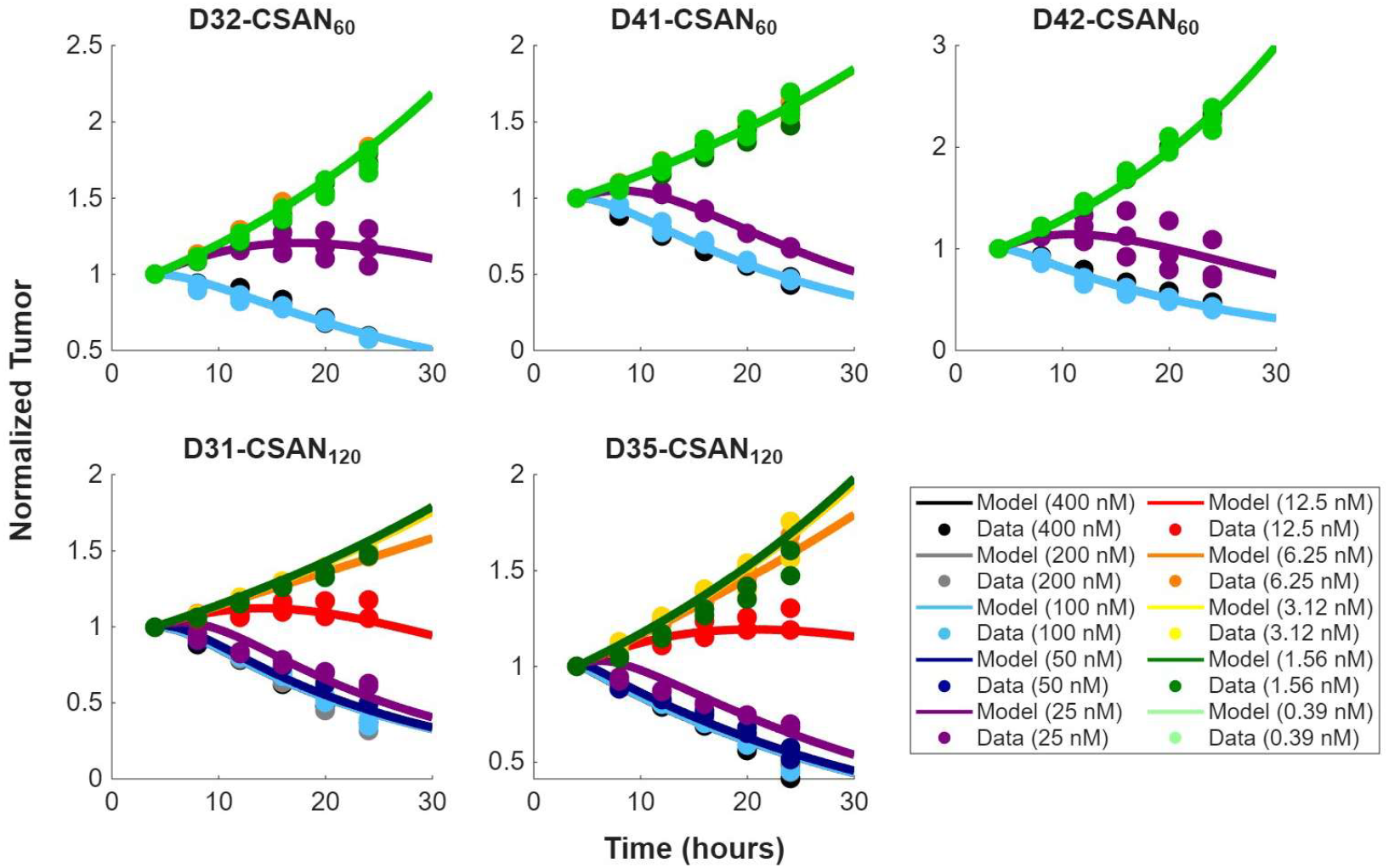
Normalized fluorescence cell counts of the A431R cell line (circular markers) in the presence of donor T cells and varying CSAN concentrations, and corresponding model predictions (eqs. S10) of A431R population dynamics (solid lines). Model parameters were fitted using donor-specific parameter estimates. (top row) Donor-specific data and model simulations for donors given CSAN_60_. (bottom row) Donor-specific data and model simulations for donors given CSAN_120_.

### Predictive modeling of TE response: validation, key drivers, and dosing predictions

We next demonstrated the robustness and predictive capabilities of the reduced model using parameter identifiability and out-of-sample validation approaches. We also identified drivers of therapeutic response variability, studied the system characteristics exhibiting donor-dependent variability, and found that CSAN efficacy is driven by the shape of dose dependence in tumor-CSAN binding dynamics. These results enable quantitative predictions of donor-specific response and minimum dosing requirements for tumor control.

#### Mechanistic inference yields identifiable model parameters and reveals sources of variability across experiments

Parameter estimation yielded unique, identifiable parameter sets whose values were highly consistent across donors under similar experimental conditions. Parameter estimates for each T cell donor are provided in Table 2. Plots of parameter values, differentiating between experimental conditions, are also shown in Figure Sl. Parameter identifiability was evaluated using the profile likelihood method (see Supplementary Profile Likelihood).

**Table 2:** Parameter values and estimates for donor-specific calibrations of eqs. S10. Parameter units are *α* (*hour*^−1^), *k*_*D*_ (*nM*), *T*_*max*_ (*hour*^−1^*) k*_*C*_ (*hour*^−1^), *n*_*T*_ (*dimensionless*), *β* ((*hour·cell*)^−1^), and *ρ* (*cells·* (*complex · hour*^−1^)). Asterisks indicate parameters with statistically significant differences in value across experimental setups (CSAN_60_ vs CSAN_120_ (see Supplementary Tumor Growth Statistics).

| Parameter | D32 | D41 | D42 | D31 | D35 |
| --- | --- | --- | --- | --- | --- |
| $\alpha$ | 0.0301 | 0.0236 | 0.0422 | 0.0225 | 0.0264 |
| $k_D$ | 60 | 60 | 60 | 120 | 120 |
| $T_{max}$ | 20 | 20 | 20 | 20 | 20 |
| $r_C$ | 0.030439 | 0.045478 | 0.046888 | 0.048627 | 0.032019 |
| $n_T^*$ | 6.3703 | 5.2191 | 5.7418 | 2.5674 | 2.6269 |
| $\beta$ | $9.4899 \times 10^{-6}$ | $9.831 \times 10^{-6}$ | $2.3217 \times 10^{-5}$ | $6.4372 \times 10^{-6}$ | $5.0321 \times 10^{-5}$ |
| $\rho^*$ | 0.50557 | 0.55559 | 0.5203 | 0.91919 | 0.73885 |

To assess whether parameter values differed systematically across the two experimental designs, two-sample t-tests were performed to compare parameters of D32, D41, and D42, with a 3:1 T cell to tumor cell ratio and 4-fold dilution increments of administered CSAN60 concentrations (nM), and those of D31 and D35, with a 5:1 T cell to tumor cell ratio and 2-fold dilution increments of administered CSAN_120_ concentrations (nM). The *p*-values are provided in Table S2 in Supplementary Estimate Analysis. Two parameters showed statistically significant differences across the two experimental designs: (i) the steepness coefficient *n*_*T*_ in the transition function *ζ* (*C*_0_), which controls the sharpness of the dose-response relationship for tumor cell conversion into the CSAN-bound state, and (ii) the T cell stimulation/proliferation proportionality constant *ρ* in eqs. S10. The two experimental regimes differ in the CSAN construct, donor cohort, E: T ratio, and dose spacing; thus, we note that it is possible that these differences can be interpreted as reflecting the overall experimental context and therefore cannot be uniquely attributed to any single factor. However, since *n*_*T*_ enters exclusively through *ζ* (*C*_0_), its difference in parameter estimates between CSAN constructs suggests that this parameter may capture CSAN-specific characteristics (e.g., effective affinity/avidity), whereas p may reflect context-dependent differences in the strength of T cell expansion and stimulation dynamics under the two experimental settings.

#### Donor-to-donor variability highlights sources of immune heterogeneity

We also assessed the variation in estimated model parameters between T cell donors, to understand the immune heterogeneity in our dataset. Coefficients of variation (COV) were calculated for each estimated parameter (Table 3). Across both experimental settings, the binding parameter between T cells and CSAN-bound tumor cells, *β*, exhibited significant variability, suggesting that effector-target cell interaction can be significantly heterogeneous between donors. In contrast, *n*_*T*_ and *ρ* showed the lowest COV values, suggesting that the CSAN-tumor binding transition dose dependence and T cell expansion processes were comparatively consistent across donor estimates within the same experimental regime.

**Table 3:** COV calculations for each of the four estimated parameters of eqs. S10, differentiated by the two distinct experimental settings (CSAN_60_ vs CSAN_120_).

| Parameter | CSAN<sub>60</sub> | CSAN<sub>120</sub> |
| --- | --- | --- |
| $r_C$ | 0.2227 | 0.2912 |
| $n_T$ | 0.0998 | 0.0162 |
| $\beta$ | 0.5521 | 1.0934 |
| $\rho$ | 0.0488 | 0.1538 |

### Predicting tumor response via leave-one-out validation

To assess the robustness, predictive power and calibration of the model, we performed a leave-one-out (LOO) validation analysis. For each donor, we systematically excluded one CSAN dose (and all associated replicates), and re-estimated the parameters using the remaining data. The resulting parameterization was then used to predict the withheld dose-response dataset. An example of donor-specific calibrated model predictions under this process is shown in Supplementary Loo Validation (Figure S3).

To quantify the predictive power of the model trained on the withheld dose-response data, we computed the ratio *R*_*i*_ to compare the sum of squared differences between (i) the single-dose predictions of the donor-specific model trained with all available data and (ii) the calibrated model with one dose excluded. Here, the subscript *i* refers to the single CSAN dose being predicted, corresponding to the dose excluded in LOO analysis.

Table 4 presents the mean and median values of *R*_*i*_ across all doses for each donor. The median values of *R*_*i*_ ≈ 1 indicate that the model has strong predictive performance when evaluate on out-of-sample doses, supporting its ability to capture generalizable relationships in the data. Across donors, mean ratios satisfy *R*_*i*_ < 1, indicating that full data calibration generally results in prediction errors slightly lower than the model calibrated on data with one dose withheld, as expected. Taken together, these results demonstrate that eqs. S10 robustly capture the essential dynamics of CSAN-mediated tumor-immune dynamics across donors without evidence of overfitting, supporting their use in predicting response to CSAN therapy.

**Table 4:**
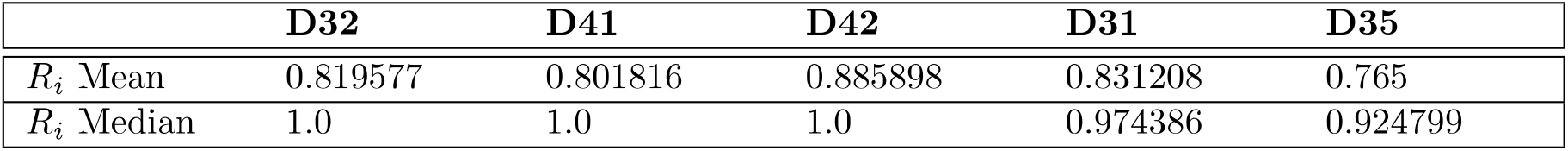
Mean and median error ratios in leave-one-out validation of model predictions, further detailed in Supplementary LOO Validation, for each donor dataset.

### Shape of dose-dependence in tumor-TE binding drives variability in early therapeutic response

In contrast to Section Donor-Specific Estimates and Section Donor Variability which characterized the donor- or context-dependent variation in inferred parameters, we next sought to quantify the relative *influence* of each parameter on variability in predicted tumor response. We employed a global sensitivity analysis via Sobol indices to quantify the influence of model parameters on tumor growth predictions, under fixed initial conditions. This variance-based approach allowed us to identify which model parameters most strongly contributed to the variation in predicted tumor quantities after 24 hours. Figure 9 provides both the first-order and total effects from this analysis.

**Figure 9:**
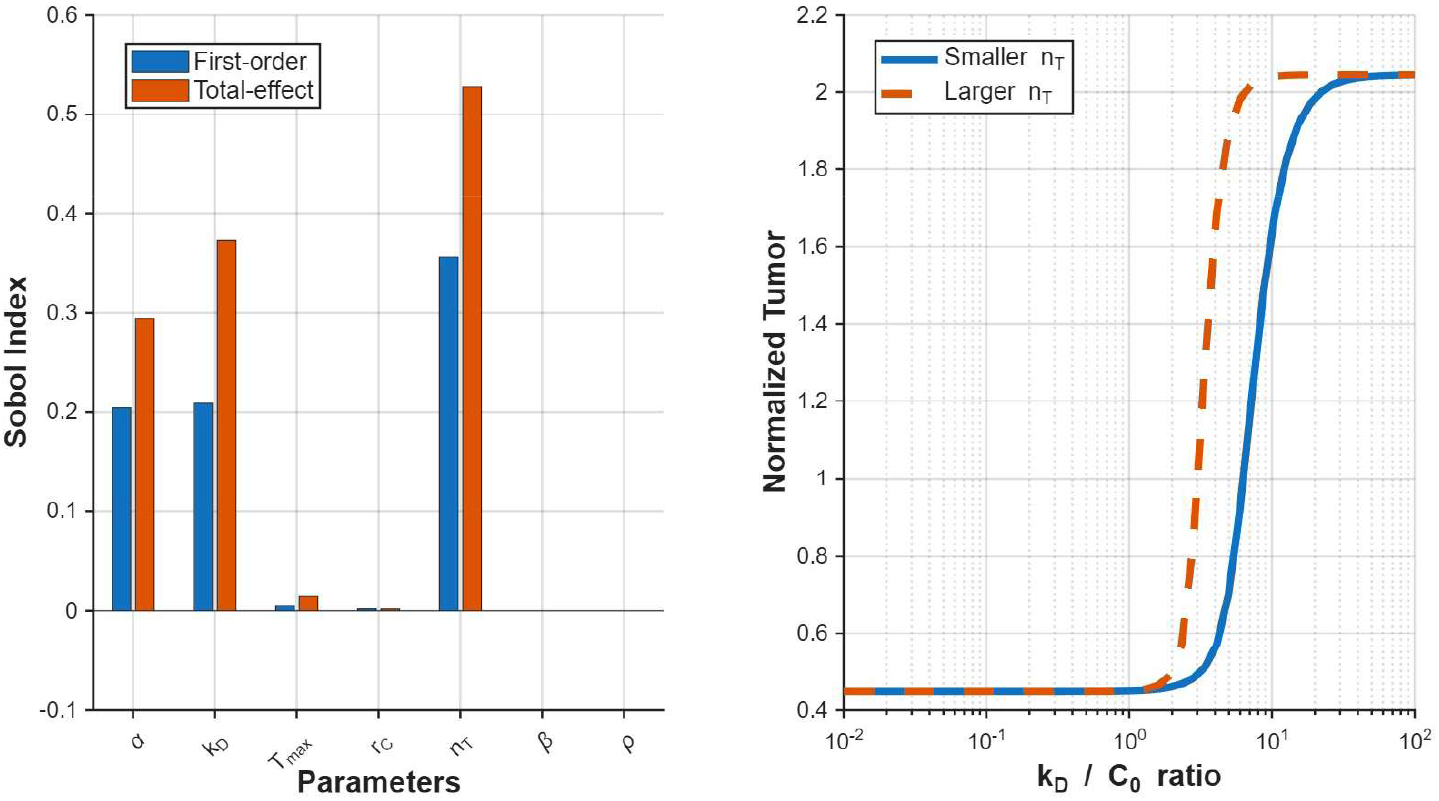
Global sensitivity analysis of eqs. S10 (left) using Sobol indices, with first-order effects in blue and total-effects in orange, quantify the relative contribution of each model parameter to the variance in the final tumor population. Parameters with higher Sobol indices exert greater influence on model output, while values near zero indicate negligible effects. Simulated, normalized tumor population dynamics (right) demonstrate the response to variation in parameter estimates, specifically for the most influential parameter identified by the Sobol indices, *n*_*T*_, under various tunable therapy conditions (*k*_*D*_ : *C*_0_). Outside of the altered *n*_*T*_, both simulation regimes were identical.

We identified three parameters that provide important contributions to the variance in model predictions at 24h: the tumor growth rate *α*, the steepness coefficient for tumor cell transitioning to the CSAN-bound state, *n*_*T*_, and the TE dissociation constant *k*_*D*_. For each influential parameter, the total effects exceeded the corresponding first-order effects, indicating that parameter interactions contributed substantially to the overall variance in model-predicted tumor populations. The importance of a is intuitive as tumor growth directly contributes to total tumor population. The parameter with the largest Sobol index was *n*_*T*_, the steepness coefficient for tumor cell transitioning to the CSAN-bound state. In eq. S10e, the parameter *n*_*T*_ governs how quickly tumor cells transition into the CSAN-bound state, provided there is a sufficient dosage of CSAN relative to the CSAN’s dissociation constant *k*_*D*_, which had the next largest index in the Sobol sensitivity analysis. Taken together, the overall process of CSAN binding and the transitioning of a tumor cell into the CSAN-bounded state heavily contributes to tumor response dynamics. Within the model (eqs. S10), trimer formation is dependent upon tumor cells having first transitioned into the CSAN-bound state. The sensitivity analysis recapitulates this notion, highlighting this process of tumor cell transitioning as a key driver of CSAN response.

#### Binding affinity-dose regime governs when donor heterogeneity influences efficacy

Given *n*_*T*_‘s heavy influence on tumor dynamics in model eqs. S10, we analytically examined how *n*_*T*_ impacts therapy response. This analysis (Supplementary Role of *n*_*T*_), demonstrates that the impact of *n*_*T*_ is dependent on the ratio of *k*_*D*_ : C_*0*_, the CSAN’s dissociation constant for the targeted tumor-associated antigen to the dosage of CSAN. For *k*_*D*_ : *C*_*0*_ < 1, any increase in *n*_*T*_ would increase the overall transitioning rate of a tumor cell into the CSAN-bound state, described by the function *ζ (C*_0_). At sufficiently large CSAN dosages, larger *n*_*T*_ improves tumor control. However, for *k*_*D*_ : *C*_0_ > 1, this transitioning rate would decrease as a result of increasing *n*_*T*_. According to the experimental data, shown in Figure 3, efficient CSAN binding, trimer formation, and tumor cell killing can be achieved at CSAN concentrations less than its respective *k*_*D*_ value (*k*_*D*_ : *C*_*0*_ > 1). This is because the Hill function within *ζ(C*_*0*_) still allows tumor cells to transition into the CSAN-bound state despite lower CSAN concentrations. At lower CSAN dosages, where *k*_*D*_ : *C*_*0*_ > 1, the transitioning rate depends on the donor-specific parameterized value of *n*_*T*_. Donors with larger estimates of *n*_*T*_ will consequently exhibit lesser tumor control compared to those with smaller estimates. In Figure 9, our model predicts that at sufficiently high CSAN concentrations, where *k*_*D*_ : *C*_*0*_ < 1, there is a sufficient level of CSAN available, so larger values of *n*_*T*_ become irrelevant. On the contrary, at lower CSAN dosages, when *k*_*D*_ : *C*_*0*_ > 1, the estimate for *n*_*T*_ begins to play a larger role. In CSAN therapies where *k*_*D*_ : *C*_*0*_ > 1, smaller estimates of *n*_*T*_ result in improved responses.

These results provide a mechanistic explanation for donor-dependent variability and yield actionable dosing insights. As both parameters in the *k*_*D*_ : *C*_*0*_ ratio are characteristics of the immunotherapy treatment, namely the CSAN used and the quantity administered, respectively, this ratio can be therapeutically tunable. Administering a CSAN dose at concentrations larger than its respective *k*_*D*_ will promote similar responses across donors, provided similar tumor growth rates, despite differences in parameter estimates for *n*_*T*_. If the CSAN dose shifts below this threshold, donor response differences become predictable with the parameter *n*_*T*_ enabling identification of donors likely to exhibit superior responses.

### Predictions on minimum dose required for tumor control

As demonstrated above, transition of a tumor cell to the CSAN-bound state plays a critical role in therapy response and tumor control. This transition rate is dependent upon three factors: the therapeutic dose (*C*_*0*_), the dissociation constant (*k*_*D*_) and the transition function’s steepness parameter (*n*_*T*_), which may depend on CSAN design. We explored the parameter space for CSAN characteristics to investigate optimal design. Figure 10 shows the necessary dose of CSANs (nM) required to elicit a 50% reduction in the tumor cell population for a given CSAN construction after 24 hours.

**Figure 10:**
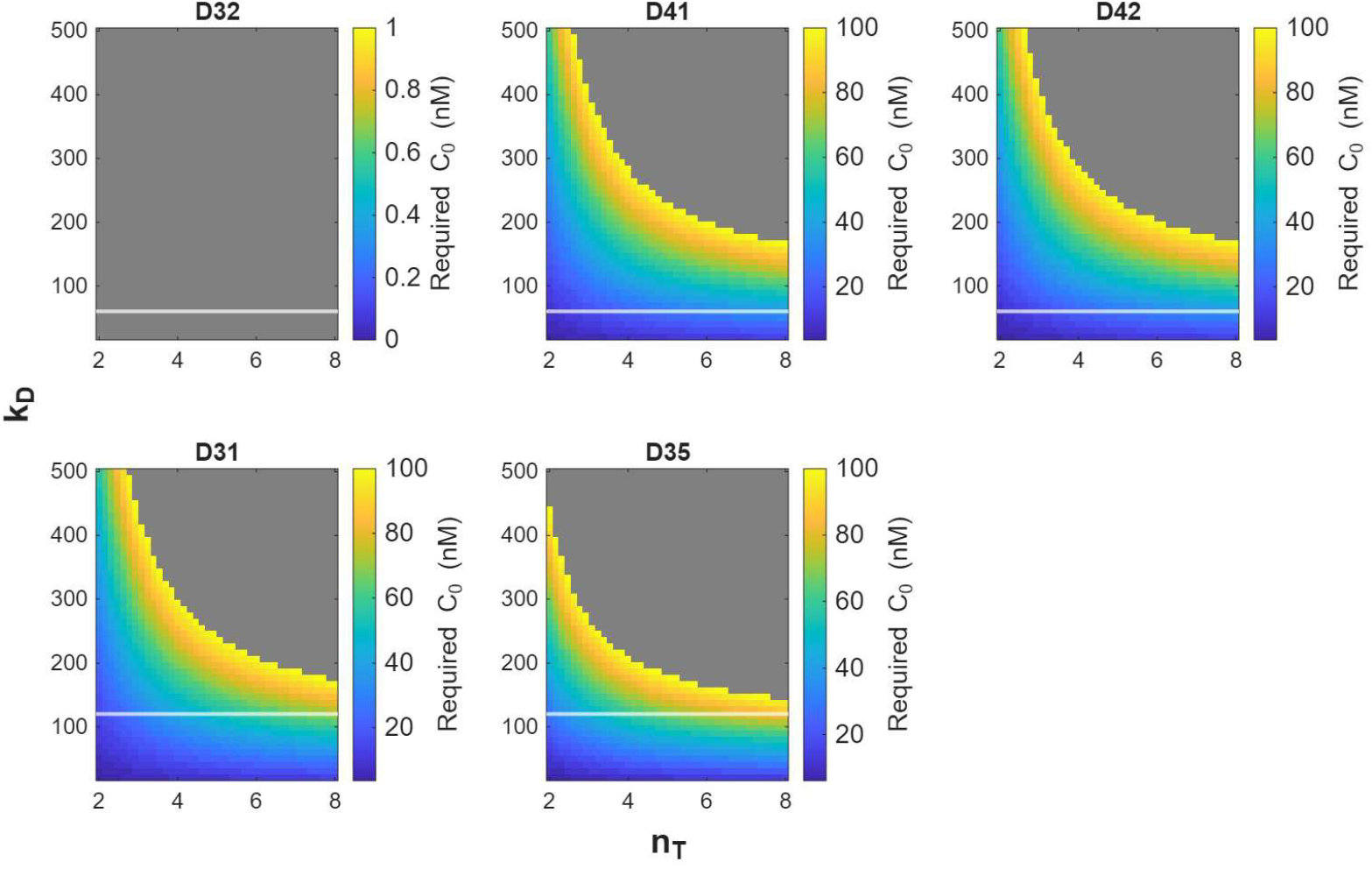
Donor-specific heatmaps indicating the model-predicted minimum dose (nM) of CSANs required for a 50% decrease in cancer cells after 24 hours. Each heatmap location corresponds to a hypothetical CSAN design parameterized by the steepness coefficient governing tumor transition to the CSAN-bound state (*n*_*T*_, X-axis) and the CSAN-tumor binding dissociation constant (*k*_*D*_, Y-axis). Colors indicate the minimum CSAN concentration (nM) required to achieve the minimum 50% reduction in tumor size within the concentration range considered (1-100 nM). Grey regions indicate parameter combinations that do achieve the 50% threshold after 24 hours with at the maximum tested dose (100 nM), i.e. would require higher concentrations. White lines indicate the kD of the CSAN used in each donor-specific dataset.

Across donors, lower values of both the CSAN-specific dissociation constant (*k*_*D*_) for the tumor-targeting antigen and the CSAN-bound transition rate’s steepness coefficient (*n*_*T*_) reduce the minimum dose concentration required to reach the threshold 50% reduction. Notably, T cells from donor D32 are unable to clear 50% of the tumor after 24 hours, regardless of the CSAN characteristics explored. This is consistent with prior parameter estimates for D32, shown in Table 2. Compared with other donor estimates, D32 exhibited the highest value of *n*_*T*_, along with the lowest cytotoxic killing rate, *r*_*C*_, and T cell expansion factor, *ρ*, all of which increase the dosing requirement for tumor control.

Finally, our inference results identified a statistically significant association between lower dissociation constants (*k*_*D*_) and larger values of the steepness coefficient (*n*_*T*_) (Table 2). These results suggest that CSAN design trade-offs may exist, where improvements in binding affinity may be offset by an increase in the steepness of the tumor binding dose-response, thus neutralizing gains in efficacy. These results highlight how both *k*_*D*_ and binding dose-response shape *n*_*T*_ jointly determine dosing requirements for efficacy, and that such trade-offs must be balanced to optimize therapeutic efficacy.

We note that the threshold of tumor clearance (50%), the maximum CSAN dose investigated (100 nM), and the 24-hour time horizon were selected for demonstration purposes; alternative choices would change the absolute dosing predictions, but the qualitative dependence on kD and *n*_*T*_ of donor-specific variability in responses would remain the same.

## Discussion

In this study, we integrated experimental measurements with mechanistic mathematical modeling to elucidate how the design features of chemically self-assembled nanorings (CSANs) and donor-specific immune variability jointly determine therapeutic efficacy. We conducted dose-response experiments using two bispecific CSAN constructs (termed CSAN_60_ and CSAN_120_) on A431R human epidermoid carcinoma cells, using primary human T cells from multiple donors across varying effector-to-tumor cell ratios. Using this data, we developed and calibrated a mathematical model that describes the coupled tumor-immune-CSAN dynamics. This framework identified key drivers of therapeutic efficacy and variability, and provides a foundation for optimizing the design and use of CSANs.

Our experiments demonstrated dose-dependent antitumor activity, with negligible killing in the absence of CSANs and variability in efficacy across T cell donors. Integrating these data with mechanistic modeling suggests that that trimer complex formation, for both CSAN constructs, relies heavily on the pathway of CSANs first binding to tumor cells, followed by subsequent binding of these CSAN-bound cells to T cells. This preference is likely related to the CSAN design, in which T cell-targeting moieties of the CSANs have a significantly larger dissociation constant, *k*_*D*_, compared to its tumor-targeting fibronectin (1000 nM to 60/120 nM, respectively) - resulting in the CSAN-bound tumor cell being the favored intermediary complex. This mechanistic asymmetry justified a model reduction that eliminated the complementary pathway (through T cell-bound intermediates) and simplified the model calibration procedure while preserving biological realism.

Model inference revealed two parameters with statistically significant differences across the two experimental regimes: the tumor-binding dose-response steepness parameter, *n*_*T*_, and the T cell stimulation/proliferation proportionality constant, *p*. In particular, the parameter *n*_*T*_ governs the steepness of the dose-dependent transition function of tumor cells into the TE-bound state, which often reflects cooperative effects in the binding process. For example, large values of *n*_*T*_ could indicate (i) a threshold in which multiple CSAN to EGFR interactions are required to form a stable immunological synapse or (ii) receptor clustering-driven affinity effects that promote multivalent binding by the same CSAN. These mechanisms can generate switch-like behavior in tumor binding, producing a steeper effective dose-response. We also observed substantial donor variability in /3, the rate of T cell binding to CSAN-bound tumor cells, suggesting that T cell engagement dynamics may be an intrinsic source of immune heterogeneity that drives patient variability.

Further analysis of *n*_*T*_ indicated that its impact on tumor dynamics is mediated through the relationship between binding affinity *k*_*D*_ and dose *C*_*0*_, When *C*_*0*_ ≳ *k*_*D*_, responses are relatively robust across donors; when *C*_*0*_ < *k*_*D*_, responses are more donor-dependent, with smaller estimated *n*_*T*_ values associated with improved tumor control. These results suggest that dosing at levels or above dissociation constant may decrease inter-patient variability in treatment response, and that at smaller doses, tumor control is still possible, and *n*_*T*_ may be a valuable mechanistic marker for predicting effectiveness.

### Design and dosing implications

We explored a large space of hypothetical CSAN characteristics and identified combinations of (*k*_*D*_, *n*_*T*_) that would minimize the dose required to achieve a given therapeutic effect. Lower *k*_*D*_ (corresponding to higher tumor affinity) and lower *n*_*T*_ (shal-lower cooperative transition in tumor-CSAN binding) together reduced the threshold dose required for reaching equivalent tumor reduction percentages. Although the dissociation constant *k*_*D*_ and Hill coefficient *n*_*T*_ are independent parameters, CSAN design features may couple them in practice. For example, increasing multivalency can lower *k*_*D*_ and stabilize binding, potentially broadening the effective therapeutic window, but may also lead to increased steepness in the binding response relationship through receptor clustering. Consistent with this hypothesis, CSAN_60_ (*k*_*D*_ = 60 nM) exhibited higher *n*_*T*_ values than CSAN120 (*k*_*D*_= 120 nM).

This hypothesis is supported by [38], whose work demonstrated that receptor clustering dynamics can be modulated by the binding affinity and valency of the interacting species. Additionally, it has been well documented that high target-binding affinities do not necessarily correspond to improved clinical outcome [39]. Furthermore, it has been reported that affinity thresholds may exist [40], above which improved binding affinities may not improve cellular functions. While this was reported for T cell functionality, this phenomenon was similarly observed by [41] for cancer vaccines. Taken together, the design choice of future multivalent TEs, like CSANs, will need to consider these relationships. Therefore, following the inferences of our current model, for the future development of optimal TE design, we believe that it is crucial to understand the mechanisms in which multivalency in immune therapeutics connects with target binding affinities and therapeutic efficacy.

The reduced model presented here preserves essential dynamics for understanding TE efficacy while achieving parameter identifiability and interpretability, and this framework can be easily generalized to other T cell engager therapies. Nevertheless, several limitations warrant consideration. The model holds the free drug concentration constant, neglecting depletion mechanisms such as internalization. Additionally, the model treats receptor binding as a Hill-type transition from an averaged unbound state to an averaged bound state, rather than a stochastic, multivalent process governing a spectrum of receptor-bound states. Our experiments span a limited time window of *in vitro* dynamics; extensions to longer-term or *in vivo* settings may reveal additional mechanisms such as immune exhaustion, cytokine release, spatial constraints and biodistribution effects. Such extensions could further enable prediction of immune-mediated toxicities (e.g. cytokine release syndrome) and evaluate mitigation strategies, including trimethoprim-mediated CSAN disassembly.

Given the potential relationship between *k*_*D*_ and *n*_*T*_, and these current limitations, a stochastic modeling perspective may be an ideal approach for future investigations. This perspective may provide a more biologically accurate depiction of TE therapies at the receptor level, perhaps helping to shed light on the relationships between physical properties of the TE and how these properties (TE design) influences therapeutic efficacy.

Overall, this work provides a quantitative foundation for understanding donor- and design-dependent heterogeneity in response to CSAN therapy, and for computationally guided preclinical optimization of CSAN design and treatment strategies. More broadly, our integrated modeling and experimental approach highlights general features of response variability that are relevant to other bispecific T cell engagers, and thus can be applied to inform the design and development of other immunotherapies.

## Supporting information

All supplemental material

