## Supplementary material for "Understanding heterogeneous responses to T cell engagers: Binding characteristics and dosing thresholds determine cytotoxic efficacy": All supplemental material

#### Experimental Methods

**Cell lines and culture conditions.** A431 human epidermoid carcinoma cells were purchased from American Type Culture Collection (ATCC, Rockville, MD) and cultured in DMEM(Gibco) supplemented with 10% heat-inactivated fetal bovine serum (FBS, Gibco) and 1X GlutaMAX (Gibco) at 37 °C with 5% CO2. These cells were modified to express the red fluorescent protein mKate2, using lentiviral transduction to generate the A431R tumor cell line. A431R cells were cultured using the same conditions as the wild-type A431 cells. Human peripheral blood mononuclear cells (PBMCs) were isolated from healthy donor blood using the Ficoll density gradient centrifugation. CD3+ T-cells were isolated from human PBMCs using the human T cell isolation kit (Akadeum Life Sciences, catalog # 13210-120) following the manufacturer's protocol. The isolated T-cells were activated and expanded using ImmunoCult human CD3/CD28 T cell activator complex (catalog # 10971) according to the manufacturer's protocol. Expanded T-cells were utilized in the cytotoxicity assays.

**Protein expression and purification.** pET28a(+) vector was used to clone the gene for the antiTCR-DHFR2, E1-DHFR2, and DHFR2-E1 fusion proteins using standard restriction digestion-based cloning. T7 Express (For E1), or Shuffle T7 Express (For anti-TCR) E. coli (New England Biolabs) were transformed using the expression plasmid. Individual clones possessing the plasmids were cultured in Lysogeny broth (LB) containing 50 $\mu$g/mL kanamycin at 37 °C. Once the optical density (OD600) reached  0.5, protein expression was induced by adding 0.5 mM isopropyl $\beta$-D-1-thiogalactopyranoside (IPTG), and the bacteria were allowed to grow for an additional 4 hours. After induction, the bacteria were pelleted and lysed, and the soluble protein from the lysate was purified using immobilized metal-affinity chromatography (IMAC) with a HisPur cobalt resin (Thermo Scientific, catalog # 89965). Purified protein was buffer exchanged into 1X PBS pH 7.4, aliquoted, and stored at -80 °C.

**Cytotoxicity assay.** On day 0, 10,000 A431R cells/100 $\mu$L in DMEM/FBS/GlutaMAX were plated in a 96-well plate and allowed to adhere overnight. Expanded T-cells were revived in RPMI 1640 medium supplemented with 10% FBS and 1X GlutaMAX. On day 1, bispecific CSANs were formed by mixing equimolar ratios of the two DHFR2 fusion proteins in the presence of 2.5 equivalents of bis-methotrexate for 1 hour at room temperature in 1X PBS. The expanded T-cells (effector cells) were counted, and 30,000 cells/75 $\mu$L in RPMI/FBS/GlutaMAX were added to the assay plate (thus giving an effector: target [E: T] ratio of 3:1). Similarly, the number of T cells were adjusted to 50,000/75 $\mu$L to achieve E: T of 5:1. 25 µL of the respective CSANs were then added to the assay plate, and the plate was transferred to a Cytation C10 live-cell confocal imager (Agilent). The red fluorescence of the A431R cells was measured every 4 hours for 24 or 48 hours, and the total red object area/well was measured. All data were normalized to $t = 0$ hours to get the kinetics of T-cell-mediated A431R cell death.

**Data processing for model calibration.** Across all experiments, fluorescence imaging was taken every 4 hours. For $\text{CSAN}_{60}$ experiments, fluorescence measurements were taken for a total of 48 hours following therapy administration. $\text{CSAN}_{120}$ experiments were imaged up to 24 hours. For consistency across dataset comparisons, time-course measurements used for model calibration were capped at 24 hours. The first measurement was taken at time $t = 0$ hours. However, few control replicates among the different donor-specific datasets showed that the initial fluorescence was larger at the initial time point than at the second ($t = 4$ hours). This unusual behavior is most likely an artifact of experimentation. To maintain consistency across datasets and to provide consistent baseline controls for model calibration, we capped the time-course data, ignoring the very first time point and started with data corresponding to time $t = 4$ hours.

#### Tumor Growth Statistics

To assess whether T cell presence significantly altered tumor growth rates, we performed two-sample $t$-tests (assuming unequal variances) comparing growth rates between tumor-only with tumor and T cell conditions for each donor-specific dataset. Growth rates were estimated by fitting exponential models to tumor fluorescence intensity data, and three replicate wells were analyzed for each condition. The null hypothesis, $H_0$, was that there is no difference in mean growth rate between the two groups. A significance level of $\alpha = 0.05$ was used for all tests. The results are summarized below:

- **D32 (A431R)**
  Tumor Growth Rates: 0.0299, 0.0344, 0.0311
  Tumor with T Cell Growth Rates: 0.0297, 0.0306, 0.0301
  Mean ± SD (Tumor Only): $0.0318 \pm 0.0024$
  Mean ± SD (Tumor + T cells): $0.0301 \pm 0.0005$
  $t$-statistic: 1.196 $p$-value: 0.2977
  **Conclusion:** No significant difference (fail to reject $H_0$)

- **D41 (A431R)**
  Tumor Growth Rates: 0.0338, 0.0351, 0.0344
  Tumor with T Cell Growth Rates: 0.0225, 0.0251, 0.0230
  Mean ± SD (Tumor Only): $0.0344 \pm 0.0007$
  Mean ± SD (Tumor + T cells): $0.0236 \pm 0.0012$
  $t$-statistic: 12.349 $p$-value: 0.0002
  **Conclusion:** Significant difference (reject $H_0$)

- **D42 (A431R)**
  Tumor Growth Rates: 0.0419, 0.0448, 0.0427
  Tumor with T Cell Growth Rates: 0.0442, 0.0413, 0.0410
  Mean ± SD (Tumor Only): $0.0431 \pm 0.0015$
  Mean ± SD (Tumor + T cells): $0.0422 \pm 0.0020$
  $t$-statistic: 0.705 $p$-value: 0.5196
  **Conclusion:** No significant difference (fail to reject $H_0$)

- **D31 (A431R)**
  Tumor Growth Rates: 0.0226, 0.0270, 0.0302
  Tumor with T Cell Growth Rates: 0.0223, 0.0247, 0.0210
  Mean ± SD (Tumor Only): $0.0266 \pm 0.0038$
  Mean ± SD (Tumor + T cells): $0.0225 \pm 0.0017$
  $t$-statistic: 1.602 $p$-value: 0.1844
  **Conclusion:** No significant difference (fail to reject $H_0$)

- **D35 (A431R)**
  Tumor Growth Rates: 0.0197, 0.0288, 0.0311
  Tumor with T Cell Growth Rates: 0.0272, 0.0268, 0.0257
  Mean ± SD (Tumor Only): $0.0265 \pm 0.0061$
  Mean ± SD (Tumor + T cells): $0.0264 \pm 0.0006$
  $t$-statistic: -0.006 $p$-value: 0.9952
  **Conclusion:** No significant difference (fail to reject $H_0$)

These results indicate a statistically significant reduction in tumor growth rate with T cell treatment only in dataset D41, while all other datasets showed no significant difference.

#### Parameters

Each parameter appearing in eqs. (1), accompanied by their corresponding units, is provided below. Accompanying each parameter is also a brief description of the biological process they represent.

Table S1: Parameters and functional terms found in eqs. (1), accompanied by brief descriptions.

| **Parameter (units)** | **Description** |
|---|---|
| $\alpha \quad (hour^{-1})$ | Exponential tumor growth rate |
| $\zeta_T(C) \quad (hour^{-1})$ | Tumor transition rate to CSAN-bound state |
| $\zeta_E(C) \quad (hour^{-1})$ | T cell transition rate to CSAN-bound state |
| $T_{max} \quad (hour^{-1})$ | Maximum transition rate of tumor cells to CSAN-bound state |
| $E_{max} \quad (hour^{-1})$ | Maximum transition rate of T cells to CSAN-bound state |
| $k_{D,T} \quad (nM)$ | CSAN:tumor cell dissociation constant |
| $k_{D,E} \quad (nM)$ | CSAN:T cell dissociation constant |
| $n_T \quad (dimensionless)$ | Hill coefficient of tumor cell transition to CSAN-bound state |
| $n_E \quad (dimensionless)$ | Hill coefficient of T cell transition to CSAN-bound state |
| $\beta \quad ((hour \cdot cell)^{-1})$ | T cell and tumor cell binding rate |
| $\beta_1 \quad ((hour \cdot cell)^{-1})$ | CSAN-bound T cell and tumor cell binding rate |
| $\beta_2 \quad ((hour \cdot cell)^{-1})$ | T cell and CSAN-bound tumor cell binding rate |
| $\beta_3 \quad ((hour \cdot cell))^{-1})$ | CSAN-bound T cell and CSAN-bound tumor cell binding rate |
| $\gamma \quad (hour^{-1})$ | Tumor-T cell dissociation rate |
| $\rho \quad (cells \cdot (complex \cdot hour)^{-1})$ | Trimer induced stimulation and proliferation rate of T cells |
| $\delta \quad (hour^{-1})$ | T cell death rate |
| $r \quad (hour^{-1})$ | Tumor cell death rate |
| $r_C \quad (hour^{-1})$ | CSAN-mediated tumor cell death rate |
| $\upsilon_T \quad (nM \cdot cell^{-1})$ | Concentration of CSANs utilized for tumor cell state transition |
| $\upsilon_E \quad (nM \cdot cell^{-1})$ | Concentration of CSANs utilized for T cell state transition |

#### Model Fitting

For each donor-specific dataset, after setting the tumor growth rate, $\alpha$, to the mean growth rates found in Section Tumor Growth Statistics and using relevant experimentally-determined values of the dissociation constants, $k_D$, calibration of the remaining parameters in Table 1 was performed using MATLAB's *fmincon*. For each dataset, all experimental data were used simultaneously (each replicate for a single nM dose of CSANs and across all CSAN dosages), resulting in a single estimated set of parameters. The least-squares error between experimental data for normalized tumor fluorescence and the total combined tumor populations from the ODE model simulation was minimized for each donor independently.

We employed several hundred initial parameter guesses randomly sampled throughout the parameter space. For each initial guess, *fmincon*'s optimization, using MATLAB's *ode15s* to handle stiff models, was computed in parallel.

For each CSAN dose, $i$, of a donor-specific dataset, we solve the system of ODEs (2), to obtain model predictions $\mathbf{x}^{(i)}(t)$ and interpolate them at the measurement time points. We define the model-predicted total tumor population ($\hat{y}^{(i)}(t_j)$) as:

$$\hat{y}^{(i)}(t_j) = x_1^{(i)}(t_j) + x_2^{(i)}(t_j) + x_4^{(i)}(t_j), \tag{S1}$$

where $x_1$, $x_2$, and $x_4$ are the corresponding state variables in the ODE model for which a tumor cell is present. The data-fitting cost for data corresponding to CSAN dose $i$ is the sum of squared errors between the model prediction and the experimental measurements. These measurements vary according to the donor-specific datasets due to differences in the number of experimental replicates. Dual replicate experiments are depicted here:

$$\text{cost}_{\text{data}}^{(i)} = \sum_{j=1}^{N_t} \left[\hat{y}^{(i)}(t_j) - y_1^{(i)}(t_j)\right]^2 + \sum_{j=1}^{N_t} \left[\hat{y}^{(i)}(t_j) - y_2^{(i)}(t_j)\right]^2, \tag{S2}$$

where $N_t$ is the number of measurement time points per dataset.

No T cell quantitative measurements were available in the experimental data. However, given the qualitative experimental observation of an approximate 2-3 fold expansion in this population, an additional penalty was added to the objective function. This penalty was proportional to the squared difference between the simulated final-to-initial T cell population ratio and the expected ratio of a 2-3 fold expansion. Specifically, first define

$$\mathrm{F}^{(i)} \equiv \frac{x_3^{(i)}(t_{\text{final}})}{x_3^{(i)}(t_1) + \varepsilon}, \tag{S3}$$

where $x_3$ is the T cell state variable of the model, $t_1$ is the initial time, $t_{\text{final}}$ is the final time point, and $\varepsilon$ avoids division by zero. The penalty term is defined as:

$$\text{penalty}^{(i)} = \begin{cases} (2 - \mathrm{F}^{(i)})^2, & \text{if } \mathrm{F}^{(i)} < 2, \\ (\mathrm{F}^{(i)} - 3)^2, & \text{if } \mathrm{F}^{(i)} > 3, \\ 0, & \text{otherwise}, \end{cases} \tag{S4}$$

and is weighted by a hyperparameter $\lambda$:

$$\text{cost}_{\text{penalty}}^{(i)} = \lambda \, \text{penalty}^{(i)}, \tag{S5}$$

with $\lambda = 0.2$. This weight was chosen so that the penalty would be comparable in scale to the summed squared error from the data-fitting cost across donor-specific datasets for previous attempts of model calibration without penalty.

The total cost used for parameter estimation is then summed over all CSAN doses of a donor-specific dataset:

$$\text{Total Cost} = \sum_{i=1}^{N_{\text{datasets}}} \left( \text{cost}_{\text{data}}^{(i)} + \text{cost}_{\text{penalty}}^{(i)} \right) \tag{S6}$$

Using this objective function, profile likelihood analysis (see Section Parameter Identifiability) was performed to assess the identifiability of parameters.

#### Nondimensionalization

The initial reduced set of model equations following simplifications (eqs. (2)) are:

$$\frac{dT}{dt} = \alpha T - \zeta(C)T, \tag{S7a}$$

$$\frac{d[CT]}{dt} = \zeta(C)T - \beta E[CT], \tag{S7b}$$

$$\frac{dE}{dt} = \rho[ECT] - \beta E[CT] + r_C[ECT], \tag{S7c}$$

$$\frac{d[ECT]}{dt} = \beta E[CT] - r_C[ECT], \tag{S7d}$$

$$\zeta(C_0) = \frac{T_{max} C_0^{n_T}}{k_D^{n_T} + C_0^{n_T}}. \tag{S7e}$$

To reduce model stiffness and to achieve efficient and appropriate parameterization, we nondimensionalize this model. Define the following relationships:

$$\hat{T} = \frac{T}{T_0}, \quad [\hat{CT}] = \frac{[CT]}{T_0}, \quad \hat{E} = \frac{E}{E_0}, \quad [E\hat{CT}] = \frac{[ECT]}{T_0}, \tag{S8}$$

where $E_0$ and $T_0$ are, respectively, the initial effector T cell and tumor cell populations. We also define a characteristic time derived from the tumor growth equation (eq. (1a)):

$$t' \equiv \alpha t$$

Across the donor-specific datasets for CSAN experiments, two sets of tumor-to-T cell ratios were used as initial conditions. To keep this nondimensionalization general and applicable to either experimental setup, we define the ratio $x \equiv E_0/T_0$.

Nondimensionalization of the left-hand side of (2) proceeds as follows (one example is presented here).

$$\frac{dT}{dt} = \frac{d}{dt}(\hat{T} \cdot T_0) = T_0 \frac{d\hat{T}}{dt} = T_0 \frac{d\hat{T}}{dt'} \frac{dt'}{dt} = T_0 \frac{d\hat{T}}{dt'} \cdot \alpha = \alpha T_0 \frac{d\hat{T}}{dt'}$$

Starting with eq. (2a), the substitutions of eqs. S8 produces an initial equation of

$$\alpha T_0 \frac{d\hat{T}}{dt'} = \alpha(\hat{T} \cdot T_0) - \zeta(C_0)(\hat{T} \cdot T_0).$$

Dividing both sides by $\alpha T_0$ and reparameterizing with the definitions $\hat{\zeta}(C_0) = \frac{\zeta(C_0)}{\alpha}$ we obtain the nondimensionalized form of eq. (2a):

$$\frac{d\hat{T}}{dt'} = (1 - \hat{\zeta}(C_0))\hat{T}.$$

The function, $\zeta(C_0)$, and its nondimensionalized form, $\hat{\zeta}(C_0)$, are independent of other cell species. For simplicity, we are not detailing its nondimensionalization here but will return to this point and show its derivation at the end.

Next, for (2b), we similarly start with

$$\alpha T_0 \frac{d[\hat{CT}]}{dt'} = \zeta(C_0)(\hat{T} \cdot T_0) - \beta \cdot (\hat{E} \cdot E_0)([\hat{CT}] \cdot T_0)$$

Division by $\alpha T_0$ and introducing $\beta^{(E)} = \frac{\beta \cdot E_0}{\alpha}$ yields

$$\frac{d[\hat{CT}]}{dt'} = \hat{\zeta}(C_0)\hat{T} - \beta^{(E)}\hat{E}[\hat{CT}]$$

For eq. (2c), we start with

$$\alpha E_0 \frac{d\hat{E}}{dt'} = \rho \cdot ([E\hat{CT}] \cdot T_0) - \beta \cdot (\hat{E} \cdot E_0)([\hat{CT}] \cdot T_0) + r_C([E\hat{CT}] \cdot T_0)$$

Dividing both sides by $\alpha E_0$ and using $\hat{\rho} = \frac{\rho}{\alpha \cdot x}$, $\beta^{(T)} = \frac{\beta \cdot T_0}{\alpha}$ and $\hat{r}_C = \frac{r_C}{\alpha \cdot x}$ yields

$$\frac{d\hat{E}}{dt'} = \hat{\rho}[E\hat{CT}] - \beta^{(T)}\hat{E}[\hat{CT}] + \hat{r}_C[E\hat{CT}]$$

Similarly for the trimer equation (2d) we start with

$$\alpha T_0 \frac{d[E\hat{CT}]}{dt'} = \beta \cdot (\hat{E} \cdot E_0)([\hat{CT}] \cdot T_0) - r_C[E\hat{CT}]T_0$$

Dividing by $\alpha T_0$ and introducing $\hat{r}_C \cdot x = \frac{r_C}{\alpha}$ yields

$$\frac{d[E\hat{CT}]}{dt'} = \beta^{(E)}\hat{E}[\hat{CT}] - x\hat{r}_C[E\hat{CT}]$$

Lastly, for equation 2e, note that $\zeta(C_0)$ is a function of the initial CSAN dosage as opposed to a dynamic state variable due to model simplifications discussed earlier. Therefore, nondimensionalization of the CSAN concentration, which is unchanging, nondimensionalizes the initial dosage by itself. As a result, this function can be expressed as:

$$\hat{\zeta}(C_0) = \frac{T_{max}}{\alpha} \cdot \frac{1}{(\frac{k_D}{C_0})^{n_T} + 1} \tag{S9}$$

The final nondimensionalized model equations are as follows:

$$\frac{d\hat{T}}{dt'} = (1 - \hat{\zeta}(C_0))\hat{T}, \tag{S10a}$$

$$\frac{d[\hat{CT}]}{dt'} = \hat{\zeta}(C_0)\hat{T} - \beta^{(E)}\hat{E}[\hat{CT}], \tag{S10b}$$

$$\frac{d\hat{E}}{dt'} = \hat{\rho}[E\hat{CT}] - \beta^{(T)}\hat{E}[\hat{CT}] + \hat{r}_C[E\hat{CT}], \tag{S10c}$$

$$\frac{d[E\hat{CT}]}{dt'} = \beta^{(E)}\hat{E}[\hat{CT}] - x\hat{r}_C[E\hat{CT}]. \tag{S10d}$$

$$\hat{\zeta}(C_0) = \frac{T_{max}}{\alpha} \cdot \frac{1}{(\frac{k_D}{C_0})^{n_T} + 1}. \tag{S10e}$$

#### Parameter Estimates

Values of eqs. (S10)'s four estimated parameters ($r_C$, $n_T$, $\beta$, and $\rho$), shown in Table 2, were plotted to visualize differences in estimates across the two experimental conditions (Figure S1. To demonstrate these differences, donor estimates corresponding to the use of $\mathrm{CSAN}_{60}$ (D32, D41, and D42) are shown with a blue circular markers while donor estimates for $\mathrm{CSAN}_{120}$ (D31 and D35) are depicted with black square markers. Using the interquartile range (IQR), not shown, of the five estimated values, we define outliers as values below or above $1.5 \cdot IQR$ from the first and third quartiles, respectively. There are no outliers for any parameter.

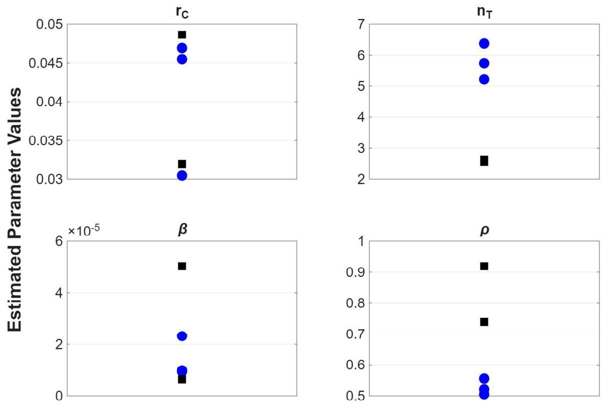


Figure S1: Plots of all estimated donor-specific parameters. Blue circles represent estimates from D32, D41, and D42 ($\mathrm{CSAN}_{60}$), while black squares indicate estimates of D31 and D35 ($\mathrm{CSAN}_{120}$).

### Parameter Identifiability

Profile likelihood analysis was used to assess parameter identifiability in the reduced model following the parameter fitting procedure. Initial analysis indicated that the parameter $T_{max}$, which represents the maximum rate of tumor cell transitioning to a CSAN-bound state, was practically unidentifiable. Specifically, below a threshold value, changes in $T_{max}$ strongly affected the model error, but above that threshold, further increases produced negligible changes in error. To address this, we fixed the parameter $T_{max}$ at a value along the lower, well-constrained edge of the profile, a value of 20 ($hour^{-1}$).

Thus, with $\alpha$ set via control experiment datasets, $k_D$ set to experimentally quantified values, and $T_{max}$ set to a constant, profile likelihood analysis was repeated for remaining parameters $n_T$, $\beta$, $\rho$, and $r_C$ and demonstrated practical identifiability. Figure S2 shows the profiles of the four remaining estimated parameters, $r_C$, $n_T$, $\beta$, and $\rho$ for the D41-calibrated model, as an example.

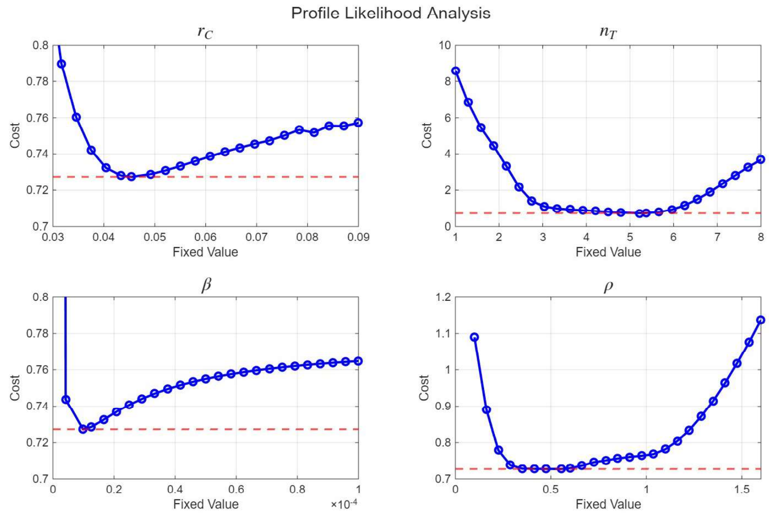


Figure S2: Profile likelihood curves for D41-estimated parameters $r_C, n_T, \beta, \rho$. Optimized parameter values, determined by the fitting procedure, yield cost function values shown as red dashed lines. Sharp troughs located on the red dashed line indicate unique parameter estimation corresponding to the minimized error.

#### Estimate Analysis

To assess whether the estimated parameters differed between the two sets of experimental conditions ($CSAN_{60}$ at 3:1 E: T vs $CSAN_{120}$ at 5:1 E: T), we performed a two-sample $t$-test for each parameter. The groups were defined as follows:

- **$CSAN_{60}$:** Datasets D32, D41, and D42
- **$CSAN_{120}$:** Datasets D31 and D35

The parameters tested included $\alpha$, the exponential growth rate of tumor cells, $r_C$, the trimer-induced tumor death rate, $n_T$, the steepness coefficient for tumor cells transitioning to a CSAN-bound state, $\beta$, the trimer formation rate, and $\rho$, the proliferation and expansion proportionality constant.

For each parameter, the MATLAB function *ttest2* was used to perform a two-sample $t$-test comparing the estimated values in $CSAN_{60}$ versus $CSAN_{120}$. This test evaluates the null hypothesis that the means of the two groups are equal. A significance threshold of $\alpha = 0.05$ was used.

Table S2: Computed $p$-values from two-sample t-tests comparing estimated model parameters between two experimental regimes (D32, D41, D42 vs. D31, D35). Asterisks indicate statistically significant differences at the 0.05 level (95% confidence interval).

| **Parameter** | **$p$-value** |
|---|---|
| $\alpha$ | 0.24 |
| $r_C$ | 0.9511 |
| $n_T$ | 0.005121 * |
| $\beta$ | 0.4734 |
| $\rho$ | 0.02286 * |

#### Leave-One-Out Validation

The "leave-one-out" method of model validation performs parameter estimation using all available data (for each donor) except one CSAN dose. Simulations of the model equations using donor-specific calibrations are then performed for the excluded CSAN dose, acting as a prediction for the donor's T cells' antitumor abilities following CSAN mediation at the excluded concentration. Error calculations between model prediction and the excluded data provide a metric for the model's predictive power. Here, we make use of the sum of squared differences. This process was repeated, evaluating predictive abilities when each dose had been excluded from the estimation process for each donor. Below are example model predictions for D31.

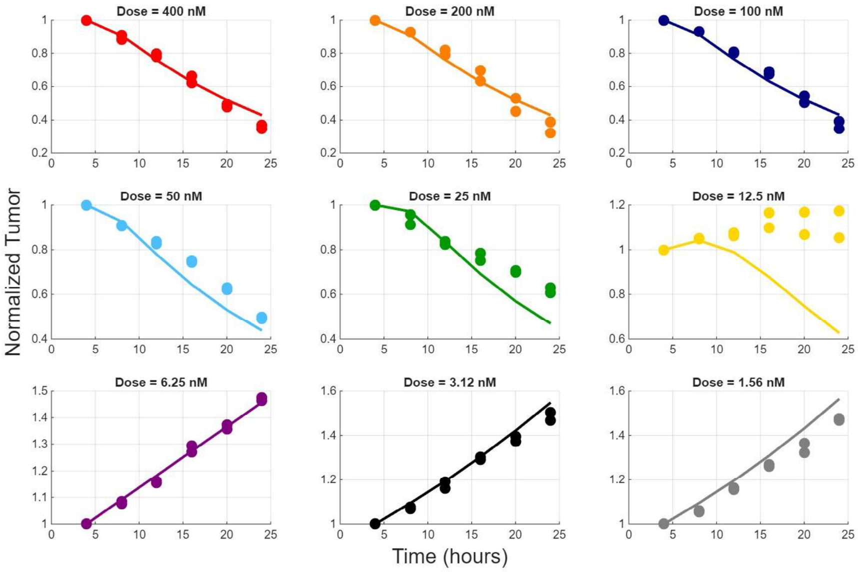


Figure S3: D31 LOO model predictions for each excluded dose of CSANs. Model predictions for a given dose are represented by solid lines whereas the excluded data used for validation is shown by solid markers.

LOO validation provides a rigorous, data-driven means of identifying whether the model captures consistent biological trends across doses or whether particular concentrations exert disproportionate influence on parameter fitting. Visual inspection of Figure S3 highlights that the model predictions were agreeable with the excluded CSAN dose data. However, one exclusion performed poorly, specifically when the CSAN dose concentration was at an intermediate level (12.5 nM for the D31 example shown above). This poor performance most likely stems from observations of the experimental data, where intermediate concentrations of CSANs (12.5 nM for D31 especially) produce larger variability in tumor fluorescence than high or low doses. This can be visually seen in Figure 3. It could be surmised that excluding data with high variability in training, therefore, contributes to poorer predictive power at doses known to exhibit high variability.

Beyond visual inspection, ratios of the sums of squared differences between the fully trained model and the LOO-calibrated model were used to quantify the relationship between the two methods of model training. This ratio is defined as:

$$R_i = \frac{E_{\text{all}}}{E_{-i}}, \tag{S11}$$

where $E_{\text{all}}$ is the fitting error obtained using all datasets, and $E_{-i}$ is the validation error when experiment $i$ is omitted during training.

Values of $R_i \approx 1$ indicate consistent model performance between full training and LOO, suggesting good generalization. Ratios $R_i < 1$ indicate that the fully calibrated model performs better when predicting the left-out experimental data (higher LOO error). This means that incorporating the excluded dataset will improve the predictive power at said concentration. $R_i > 1$ may indicate that the omitted dataset is easier to predict than the others.

This ratio provides insight into the model's stability and predictive capability. Consistently minor deviations between $E_{\text{all}}$ and $E_{-i}$ (i.e., $R_i \approx 1$ imply that the model captures the underlying biological or physical mechanisms common to all datasets. In contrast, large deviations or outliers in these metrics may reveal experiment-specific effects, model structural limitations, or overfitting to particular datasets.

#### Analytical Role of $n_T$

We define the nondimensional total tumor mass

$$S(t') \;=\; \hat{T}(t') + [\hat{CT}](t') + [\hat{ECT}](t').$$

Using the ODEs (S10a)–(S10d) we form the time derivative of $S$:

$$\begin{aligned}\frac{dS}{dt'} &= \frac{d\hat{T}}{dt'} + \frac{d[\hat{CT}]}{dt'} + \frac{d[\hat{ECT}]}{dt'} \\ &= \big(1 - \hat{\zeta}(C_0)\big)\hat{T} + \hat{\zeta}(C_0)\hat{T} - \beta^{(E)}\hat{E}\,[\hat{CT}] + \beta^{(E)}\hat{E}\,[\hat{CT}] - x\hat{r}_C[\hat{ECT}] \\ &= \hat{T} - x\hat{r}_C\,[\hat{ECT}]. \end{aligned} \tag{S12}$$

Although $\frac{dS}{dt'}$ does not contain $\hat{\zeta}(C_0)$ explicitly, the state $[\hat{ECT}](t')$ evolves under dynamics that do depend on $\hat{\zeta}(C_0)$. Hence $n_T$ affects $S$ only *indirectly* by changing the partitioning of mass between $\hat{T}, [\hat{CT}]$ and $[\hat{ECT}]$ through $\hat{\zeta}(C_0)$.

**Sensitivity of $\hat{\zeta}(C_0)$ to $n_T$.** The transitioning function is given by:

$$\hat{\zeta}(C_0) = \frac{T_{\max}}{\alpha} \cdot \frac{1}{\left(\frac{k_D}{C_0}\right)^{n_T} + 1}.$$

We differentiate $\hat{\zeta}(C_0)$ with respect to $n_T$ to see how increasing the Hill exponent affects the transition rate:

$$\begin{aligned}\frac{\partial\hat{\zeta}(C_0)}{\partial n_T} &= \frac{T_{\max}}{\alpha} \cdot \frac{\partial}{\partial n_T}\left[\frac{1}{\left(\frac{k_D}{C_0}\right)^{n_T} + 1}\right] \\ &= \frac{T_{\max}}{\alpha}\left(-\frac{\left(\frac{k_D}{C_0}\right)^{n_T}\ln\!\left(\frac{k_D}{C_0}\right)}{\left(\left(\frac{k_D}{C_0}\right)^{n_T} + 1\right)^2}\right). \end{aligned} \tag{S13}$$

The sign of $\partial\hat{\zeta}(C_0)/\partial n_T$ is therefore determined by $\ln\!\left(\frac{k_D}{C_0}\right)$:

- If $\dfrac{k_D}{C_0} > 1$ (i.e. $k_D > C_0$), then $\ln\!\left(\frac{k_D}{C_0}\right) > 0$ and $\dfrac{\partial\hat{\zeta}(C_0)}{\partial n_T} < 0$. Increasing $n_T$ *decreases* $\hat{\zeta}(C_0)$.
- If $\dfrac{k_D}{C_0} < 1$ (i.e. $k_D < C_0$), then $\ln\!\left(\frac{k_D}{C_0}\right) < 0$ and $\dfrac{\partial\hat{\zeta}(C_0)}{\partial n_T} > 0$. Increasing $n_T$ *increases* $\hat{\zeta}(C_0)$.
- If $k_D = C_0$, the derivative is zero and $\hat{\zeta}(C_0)$ is locally insensitive to $n_T$.

Thus, the effect of raising the Hill coefficient $n_T$ depends qualitatively on the ratio $k_D/C_0$: raising $n_T$ sharpens the "switch" and pushes $\hat{\zeta}(C_0)$ toward 0 or 1 depending on whether the baseline point lies on the low or high side of the switch.
